# Wounding and insect feeding trigger two independent MAPK pathways with distinct regulation and kinetics

**DOI:** 10.1101/855098

**Authors:** Cécile Sözen, Sebastian T. Schenk, Marie Boudsocq, Camille Chardin, Marilia Almeida-Trapp, Anne Krapp, Heribert Hirt, Axel Mithöfer, Jean Colcombet

## Abstract

Wounding is caused by abiotic and biotic factors and triggers complex short- and long-term responses at the local and systemic level. These responses are under the control of complex signaling pathways, which are still poorly understood. Here, we show that the rapid activation of MKK4/5-MPK3/6 by wounding is independent of jasmonic acid (JA) signaling and that, contrary to what happens in tobacco, this fast module does not control wound-triggered JA accumulation in Arabidopsis. We also demonstrate that a second MAPK module, constituted by MKK3 and the clade-C MAPKs MPK1/2/7, is activated by wounding in an independent manner. We provide evidence that the activation of this MKK3-MPK1/2/7 module occurs mainly through wound-induced JA production via the transcriptional regulation of upstream clade-III MAP3Ks and particularly MAP3K14. We show that *mkk3* mutant plants are more susceptible to the larvae of the generalist lepidopteran herbivore *Spodoptera littoralis*, indicating that the MKK3-MPK1/2/7 module is involved in counteracting insect feeding.

**One sentence summary:** Wounding induces the parallel activation of a rapid signaling module (MKK4/5-MPK3/6) and a JA-dependent slow one (MAP3K14-MKK3-MPK1/2/7/14) to restrict insect feeding.

## Introduction

Wounding is a common stress for plants that can be caused by abiotic factors such as wind, heavy rain, hail and snow or during biotic interactions, mostly with herbivorous organisms such as insects. Injury may cause harsh damages to plant tissues and facilitate the entry of pathogens (Savatin et al., 2014). Plants respond to these challenges by activating several mechanisms to rapidly heal tissues and restrict potential pathogen entry. The cuticle and trichomes are important constitutive structures involved in the prevention of wounding. Once wounding occurred, intracellular molecules released from dead cells or damaged cell wall components act as signaling molecules named DAMP (for Damage Associated Molecular Pattern) (Maffei et al., 2012). Additionally, in the vicinity of wound sites, living cells sense the mechanical disturbance through the activation of mechanosensitive channels which trigger intracellular signaling pathways and local responses (Farmer et al., 2014). Such responses are mediated by efficient and complex intracellular signaling mechanisms involving phosphorylation, lipid, ROS (Reactive Oxygen Species) and Ca^2+^ signaling, and the production of phytohormones leading to important gene expression reprogramming and long distance signaling (Savatin et al., 2014). Among the phytohormones, jasmonates, members of the oxylipin family, seem to play an overriding role (Wasternack and Hause, 2013). Jasmonates and in particular JA are produced very rapidly by herbivory-induced wounding and regulate a large number of wound-induced genes (Reymond et al., 2004). The isoleucine conjugate of JA, (+)-7-*iso*-jasmonoyl-L-isoleucine (JA-Ile), is the bioactive form of jasmonates (Fonseca et al., 2009).

Mitogen-Activated Protein Kinase (MAPK) modules are very well conserved signaling pathways found in all eukaryotes (Colcombet and Hirt, 2008). A MAPK module is minimally constituted of three kinases, a MAP3K (or MAP2K Kinase), a MAP2K (or MAPK Kinase) and a MAPK, which are able to phosphorylate and thereby activate each other sequentially. These kinases are encoded by large gene families, for which we have incomplete functional information (Colcombet and Hirt, 2008). Since their discovery in plants several decades ago, a number of the MAPK components have been involved in biotic and abiotic stress signal transductions as well as developmental processes (for review (Xu and Zhang, 2015; Suarez-Rodriguez et al., 2010)). They have notably been shown to function at an early step of wound signaling. For example, alfalfa MMK4 and tobacco WIPK are activated within minutes by leaf wounding (Bogre et al., 1997; Seo et al., 1999). Generally, after a peak at 15 minutes, their activities decline to basal levels within one hour after wounding (Bogre et al., 1997; Seo et al., 1995, 1999; Usami et al., 1995). In Arabidopsis, wounding was first shown to activate both MPK4 and MPK6 (Ichimura et al., 2000). More recently, a more complete module, namely MKK4/5-MPK3/6, was identified to stimulate ethylene production in response to wounding (Li et al., 2017). Interestingly, several early studies suggested that this MAPK activation also controls JA production (Ahmad et al., 2016). Indeed, tobacco lines in which WIPK has been either silenced or overexpressed have lower and higher JA amounts after wounding, respectively (Seo et al., 2007, 1999, 1995). Additionally, JA was also shown to be a modulator of MPK6 activity, suggesting a feedback loop to fine-tune JA homeostasis (Takahashi et al., 2007). These MAPKs belong to clades A and B, which code for the well characterized iconic “stress-activated MAPKs”, but data also suggest that less studied members of the family also transduce wound signals. For example, MPK8, a clade-D MAPK, as well as the clade-C MAPKs MPK1/2 were shown to be activated by wounding (Takahashi et al., 2011; Ortiz-Masia et al., 2007).

*Arabidopsis* MPK1 and MPK2, together with MPK7 and MPK14, define the clade-C MAPKs, which were shown to function downstream of the atypical MAP2K MKK3 (Colcombet et al., 2016). The MKK3-MPK7 module was for example reported to play a role during plant–pathogen interactions as well as during drought perception; in particular, MPK1 is activated by ROS produced during the interaction with pathogenic bacteria and the drought-induced phytohormone abscisic acid (ABA) (Danquah et al., 2015; Dóczi et al., 2007). Coherently, an *mkk3* mutant is hypersensitive to infection by *Pseudomonas syringae* DC3000 and also more sensitive to water deficit. In the context of ABA signaling, the MKK3-MPK1 module is activated by the clade-III MAP3Ks MAP3K17 and MAP3K18 through transcriptional regulation which induces a delay in the activation of the module (Danquah et al., 2015; Boudsocq et al., 2015). In this present study, we characterized the MAPK-dependent phosphorylation cascades in response to tissue lesions. We unveil the coexistence of two independent MAPK modules, which are activated by wounding with different kinetics. We also revisited the knowledge about the JA-MAPK relationship and provide evidence that JA production is not controlled by stress-activated iconic MAPKs in Arabidopsis.

## Results

### Clade-C MAPK activation by wounding strictly depends on MKK3

Some clade-C MAPKs were previously shown to be activated by wounding and to act downstream of MKK3 in response to H_2_O_2_ and ABA (Danquah et al., 2014; Ortiz-Masia et al., 2007; Dóczi et al., 2007). To investigate whether MPK2 also acts downstream of MKK3 in the wounding response, MPK2 was immunoprecipitated using a specific antibody from wounded leaves of WT and *mkk3* KO plants and its activity assayed as the ability to phosphorylate the heterologous substrate Myelin Basic Protein (MBP). As clade-C activation by ABA occurs slowly, we performed a long kinetics up to 4 hours (Danquah et al., 2015). In Col-0 plants, MPK2 was activated from 30 minutes to 2 hours after wounding (fig. 1A). This activation was totally abolished in *mkk3-1* and *mkk3-2* mutants (fig. 1A and S1A) and recovered in *mkk3-1* plants transformed with the whole genomic *MKK3* locus C-terminally fused to *Yellow Fluorescent Protein* (*YFP*) (*mkk3-1 MKK3-YFP*) (fig. S1B). Because the MPK2-specific antibody is unable to detect MPK2 protein in western blot (Ortiz-Masia et al., 2007), we cannot exclude that the increase of MPK2 activity in response to wounding is due to a wound-dependent accumulation of MPK2. To challenge this hypothesis, plants expressing the *MPK2* locus C-terminally fused to a *Human influenza hemagglutinin* (*HA*) sequence were generated in both WT and *mkk3-1* genetic backgrounds. HA-immunoprecipitation from WT, but not from *mkk3-1* wounded leaves, revealed a transient increase of MPK2-HA activity with a similar kinetics as endogenous MPK2, and without any variation in protein amounts (fig. 1B). MPK2 together with MPK1, MPK7 and MPK14 belong to the clade-C MAPKs. In protoplasts, MPK1/2/7/14 seem to function in a similar module downstream of MKK3 (Danquah et al., 2015). To monitor if other clade-C MAPKs are activated by wounding, plants expressing the *MPK1* and *MPK7* loci fused to an *HA* tag were generated and subjected to wounding. Both MPK1-HA and MPK7-HA were transiently activated by wounding following the same kinetics as MPK2 (fig. S2A and S2B). MPK7-HA activation was also dependent on MKK3 as it did not occur in the *mkk3-1 MPK7-HA* background (fig. S2B). In addition, MPK1-HA and MPK7-HA proteins amounts were unaffected by wounding (fig. S2A and S2B). Altogether, these results demonstrate that clade-C MAPKs are activated by wounding in an MKK3-dependent manner with a kinetics that is considerably slower when compared to flg22-induced activation of the well-studied MPK3/6, which occurs in less than 2 minutes with a peak around 10-15 minutes (fig. S3) (Ranf et al., 2011).

**Fig. 1.**
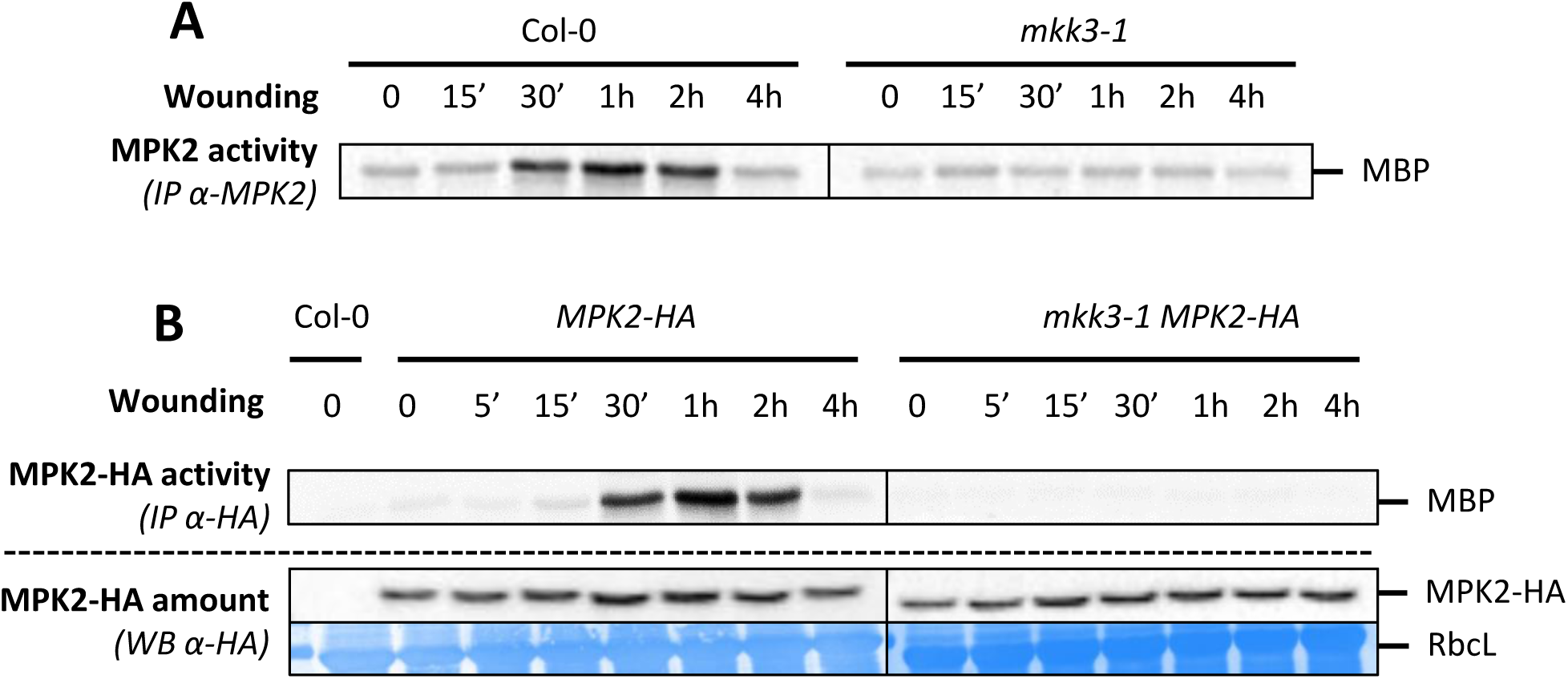
MPK2 activation by wounding depends on MKK3. **A.** Kinase activity of MPK2 after immunoprecipitation with an anti-MPK2 specific antibody from Col-0 and *mkk3-1* leaves following wounding at the indicated times. **B.** Kinase activity of MPK2 after immunoprecipitation with an anti-HA antibody from leaves of plants expressing an HA-tagged version of MPK2 in Col-0 and *mkk3-1* leaves following wounding at the indicated times. Protein amount is monitored by western-blot using anti-HA antibody. Equal loading is controlled by Coomassie staining.

### MKK3 and clade-III MAP3Ks interact and constitute a functional module when expressed in Arabidopsis mesophyll protoplasts

We previously formulated the hypothesis that the MKK3-MPK1/2/7/14 module functions downstream of clade-III MAP3Ks, namely MAP3K13-20 (Colcombet et al., 2016). MKK3 was indeed shown to interact in Yeast 2-hybrid (Y2H) assays with MAP3K15-20 but with none of the 8 tested clade-I and clade-II MAP3Ks (fig. S4). However, MKK3 did not interact neither with MAP3K13 and MAP3K14 that are the only clade-III MAP3Ks predicted to possess transmembrane domains at their carboxyl-termini (Schwacke et al., 2003). As transmembrane domains may trigger Y2H false negatives, we generated truncations of Activation Domain (AD)-MAP3K13 and AD-MAP3K14 fusions missing their 143 and 116 C-terminal amino acids, respectively, corresponding to the putative transmembrane domains. Indeed, both truncated forms of MAP3K13 and MAP3K14 were able to interact with MKK3 in Y2H (fig. 2A).

**Fig. 2.**
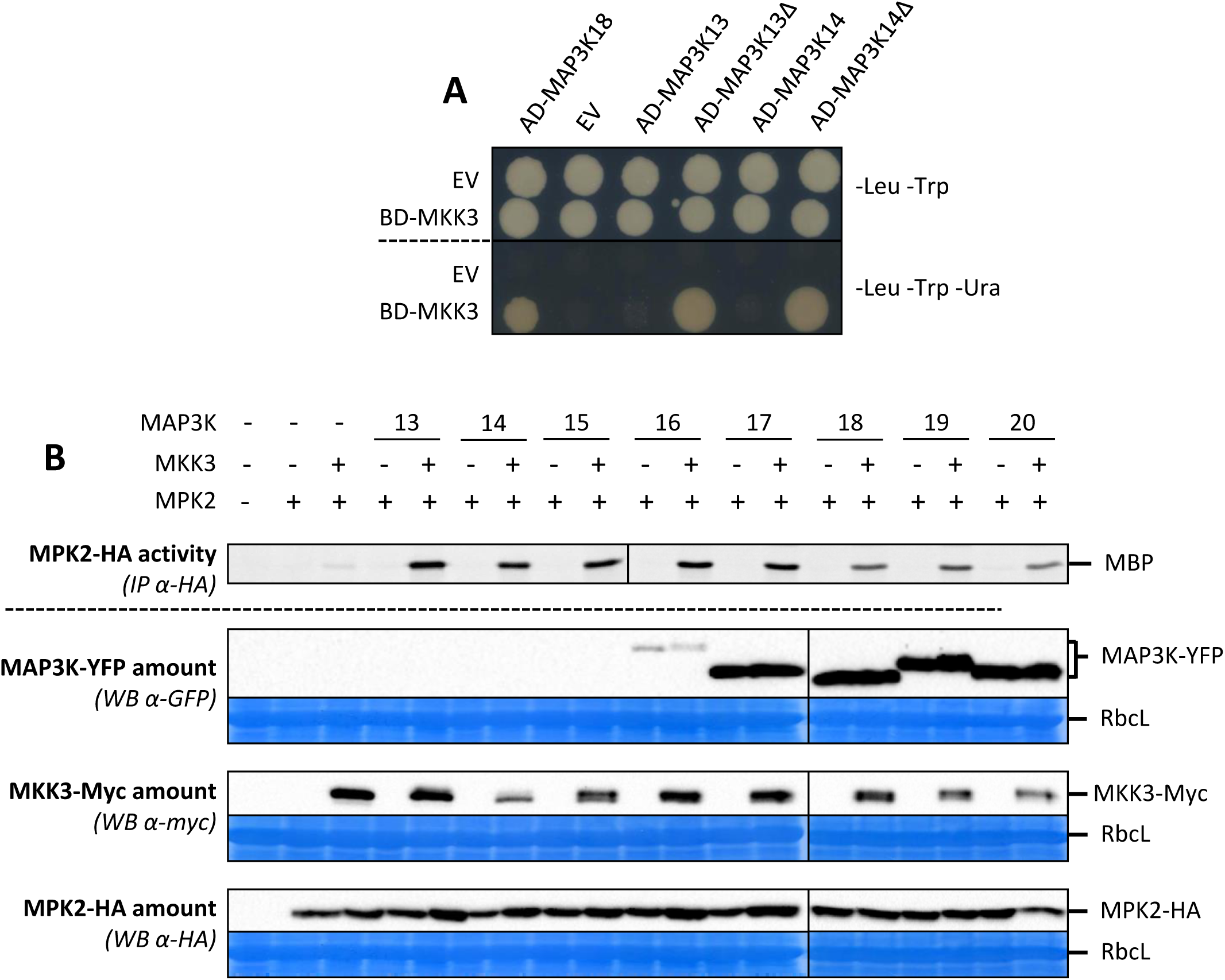
Functional reconstitution of MKK3 modules. A. Yeast two-hybrid analysis of the interaction between MKK3 and WT and C-terminal truncated forms (Δ) of MAP3K13 and MAP3K14. EV stands for Empty Vector. B. Kinase activity of HA-immunoprecipitated MPK2 transiently expressed in *mkk3-1* mesophyll protoplasts in the presence or absence of MKK3 and clade-III MAP3Ks. Western-blots show protein expression levels. Equal loading is controlled by Coomassie staining.

To test whether these MKK3-interacting MAP3Ks are able to activate MKK3-MPK2 *in planta*, we took advantage of the transient expression system in Arabidopsis mesophyll protoplasts. YFP-tagged versions of the 8 clade-III MAP3Ks were co-expressed with MPK2-HA in protoplasts generated from *mkk3-1* leaves in the absence or presence of MKK3-Myc (fig. 2B). The activity of MPK2 was assayed after immunoprecipitation with anti-HA antibodies. MPK2-HA was activated only when both MKK3 and one of the tested MAP3Ks were co-expressed together. Surprisingly, MAP3K13/14/15/16-YFP proteins were not or barely detectable, despite their ability to activate MPK2 in an MKK3-dependent way. In the case of MAP3K14, we confirmed that the module activation is dependent on kinase activity as MAP3K14^D140A^, mutated in its active site, could not activate the MKK3-MPK2 module (fig S5). Overall, these results indicate that clade-III MAP3Ks-MKK3-MPK2 and other clade-C MAPKs form functional modules *in planta*.

### Wounding induces clade-III MAP3K transcriptional regulation

We have shown that the slow ABA-induced activation of MPK7, a close homolog of MPK2, was dependent on *de novo* protein synthesis (Danquah et al., 2015). To check whether a similar mechanism is involved in the wound-induced activation of MPK2, the protein biosynthesis inhibitor cycloheximide (CHX; 100 µM) was sprayed onto plants expressing HA-tagged versions of MAPKs prior to wounding. After immunoprecipitation and kinase assays, MPK1/2/7-HA were not activated by wounding anymore, whereas the protein amounts of the MAPKs were unchanged (fig. 3A, S6A and S6B). Additionally, MKK3-YFP levels were only weakly affected by spraying CHX onto *mkk3-1 MKK3-YFP* plants prior to wounding (fig. 3B). These results suggest that the wound-induced activation of clade-C MAPKs requires the *de novo* synthesis of proteins acting upstream of MKK3. Based on these results, we hypothesized that transcriptional regulation could be a conserved regulation feature of clade-III MAP3Ks and could be a requirement to activate clade-C MAPKs. Using RT-qPCR, expression of clade-III *MAP3K* genes in WT leaves was analyzed in a wounding time-course (fig. 3C). We observed that *MAP3K14* transcript levels highly increased from 15 minutes post-wounding with a peak at 30 minutes. *MAP3K15/17/18*/*19* were also induced, but later and at more moderate levels. Additionally, using *map3k17map3k18* plants complemented with a *YFP*-tagged *MAP3K18* locus (Danquah et al., 2015), MAP3K18-YFP protein accumulated with a corresponding slow kinetics (fig. S7). Overall, these expression data are compatible with the slow MPK2 activation by wounding and suggest that MAP3K14 and other clade-III MAP3Ks could play a role in the wound-triggered MKK3-MPK1/2/7/14 activation.

**Fig. 3.**
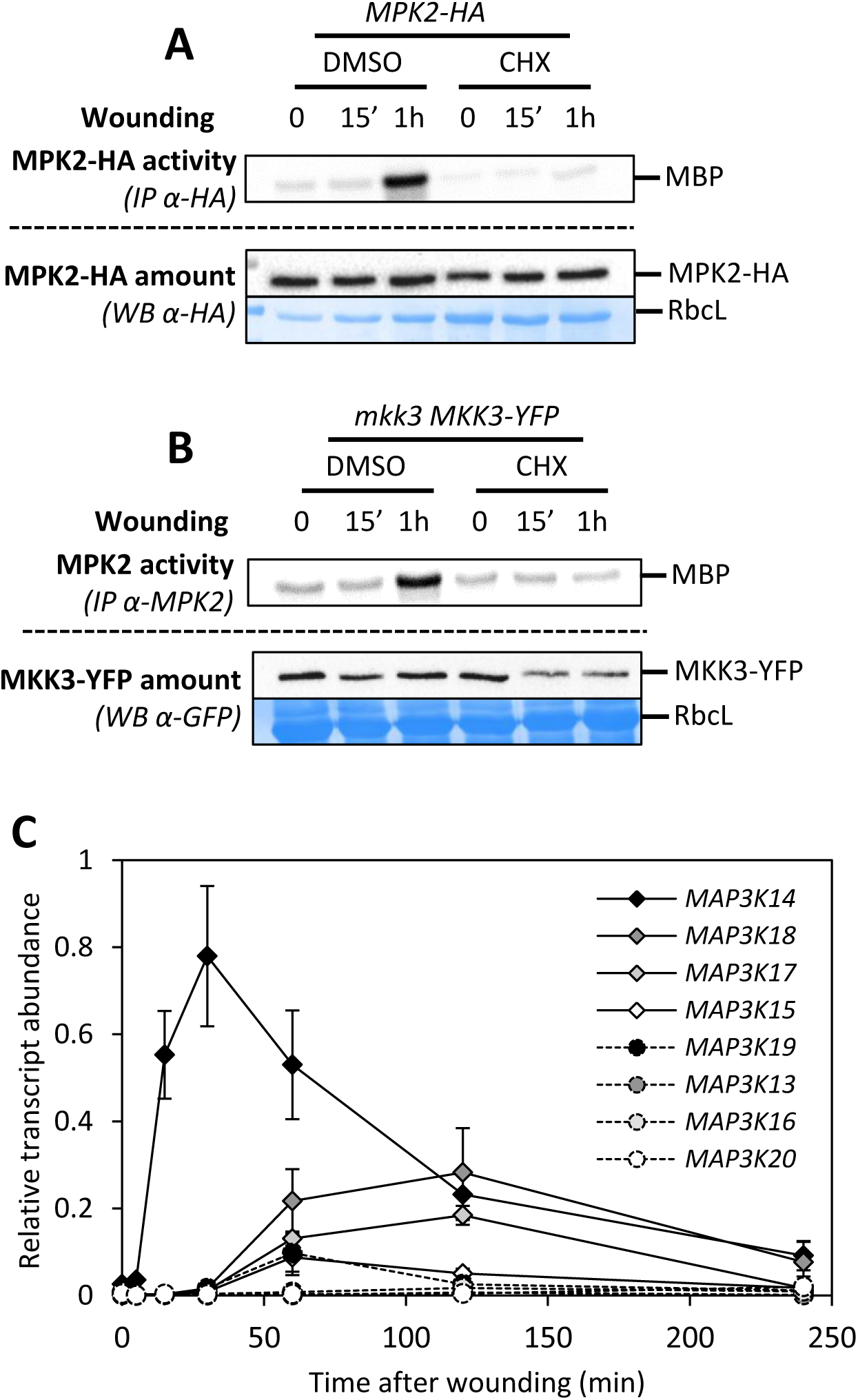
MPK2 activation by wounding requires protein synthesis. **A and B.** Kinase activity of MPK2 after immunoprecipitation with anti-HA (A) and anti-MPK2 (B) antibodies from leaves of indicated genetic background following 100µM CHX and MOCK (DMSO) spraying prior to wounding. Protein amount is monitored by western-blot using anti-HA (A) and anti-GFP (B) antibodies. Equal loading is controlled by Coomassie staining. **C.** qRT-PCR analysis of clade-III *MAP3K* genes in response to wounding. Transcript levels are expressed relative to *ACTIN2* reference gene. Values are mean±SE of 3 biological replicates.

### MAP3K14 contribute to MPK2 activation by wounding

To obtain genetic evidence that *MAP3K14* plays a role in wound-dependent MKK3 module activation, we first identified plants carrying a homozygous T-DNA insertion in the *MAPK14* gene (fig. S8A). Surprisingly, in this background, MPK2 activation was higher than in WT plants (fig. S8B). We then realized that the T-DNA insertion at position 1041 after the start codon was predicted to truncate the C-terminal tail without affecting the kinase domain. Expression of the predicted truncated protein MAP3K14-1 in *mkk3-1* mesophyll protoplasts was indeed able to activate the MKK3-MPK2 module (fig. S8C). Importantly, whereas MAP3K14 was not detected in western-blots, MAP3K14-1 showed a strong accumulation, potentially explaining the MPK2 over-activation in *map3k14-1* and suggesting that the MAP3K14 C-terminus could contain a regulatory domain for protein stability. In order to consolidate the gain-of-function role of MAP3K14-1, we used CRISPR/Cas9 technology to create two loss-of-function lines, referred to as *map3k14-CR1* and *map3k14*-*CR2*, which show a single base pair insertion (A and T, respectively) at position 550 after the start codon. The resulting frame shift generates an early stop codon leading to a truncation in the MAP3K14 kinase domain (fig. 4A). *map3k14-CR1* and –CR2 plants subjected to wounding showed a reduction of MPK2 activation particularly at 30 min (fig. 4B and fig. S9). This result is in agreement with *MAP3K14* being the only clade-III *MAP3K* transcriptionally induced at early time points after wounding (fig. 3C). Overall, these data support the hypothesis that MAP3K14 initiates the wound-activation of the MKK3-MPK2 module but also indicates that other MAP3Ks take over MPK2 activation at later time points.

**Fig. 4.**
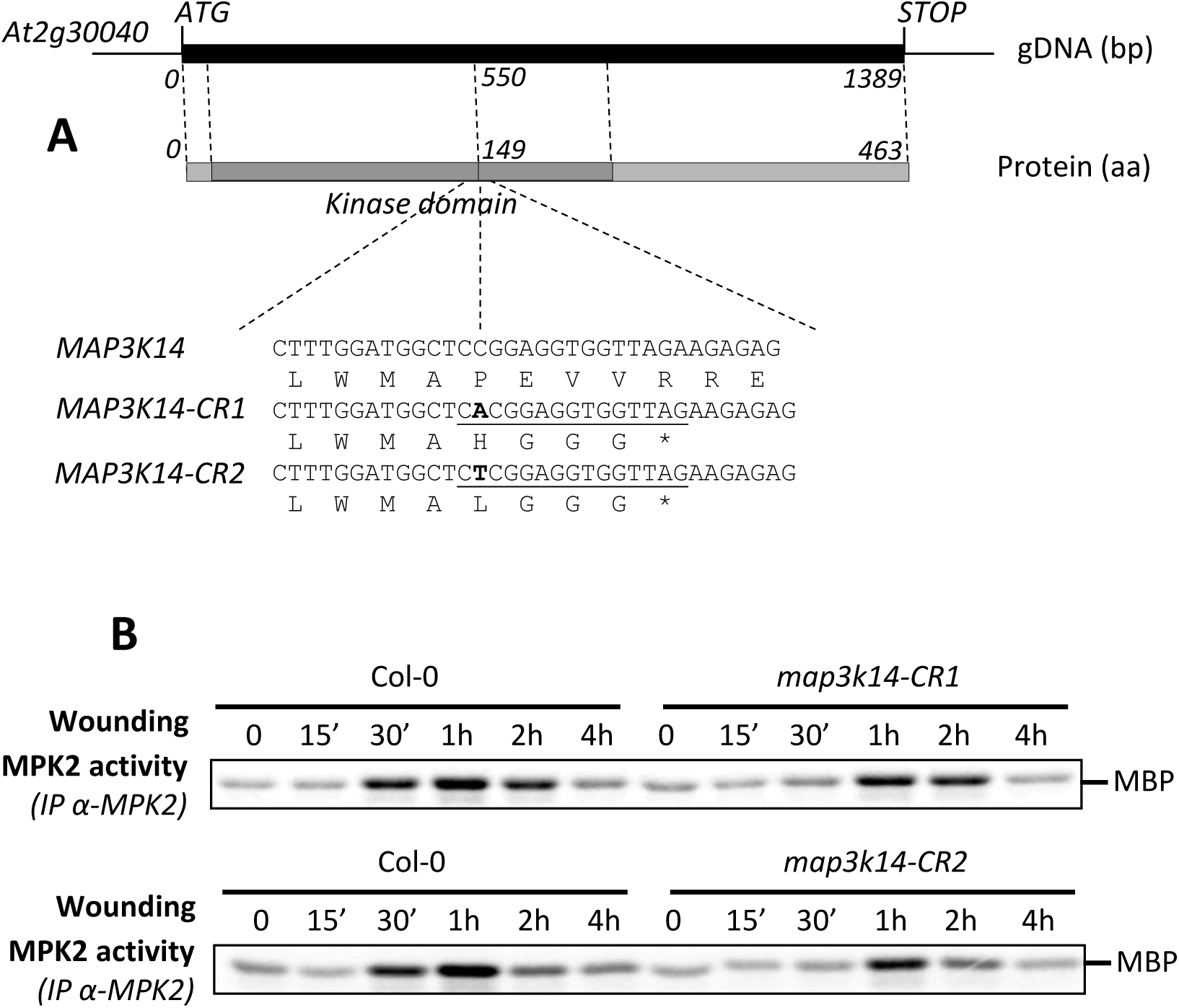
MAP3K14 plays a role in the activation of MPK2 by wounding. **A.** Genomic structure of *MAP3K14* and *CRISPR* lines used in this work. **B.** Kinase activity of MPK2 after immunoprecipitation with an anti-MPK2 antibody from WT and *map3k14-CR1* and *-CR2* leaves following wounding.

### Jasmonic acid regulates wound-triggered MKK3 module activation

As shown in Figure 5A, Jasmonic acid (JA) is an important phytohormone that is rapidly produced upon wounding in an MKK3-independent manner, as well as its isoleucine conjugate, JA-Ile, and its biosynthetic precursor, cis-OPDA. We further investigated whether JA plays a role in the wound-triggered MAPK activation. Upon JA treatment (50 µM) of 10-day-old plantlets, the activity of endogenous MPK2 increased with a rather slow kinetics (peaking at 30-60 min depending on the experiments) in WT but not in *mkk3-1* (fig. 5B). Similarly, JA activated MPK2-HA, MPK1-HA and MPK7-HA without affecting protein amounts (fig. S10).

**Fig. 5.**
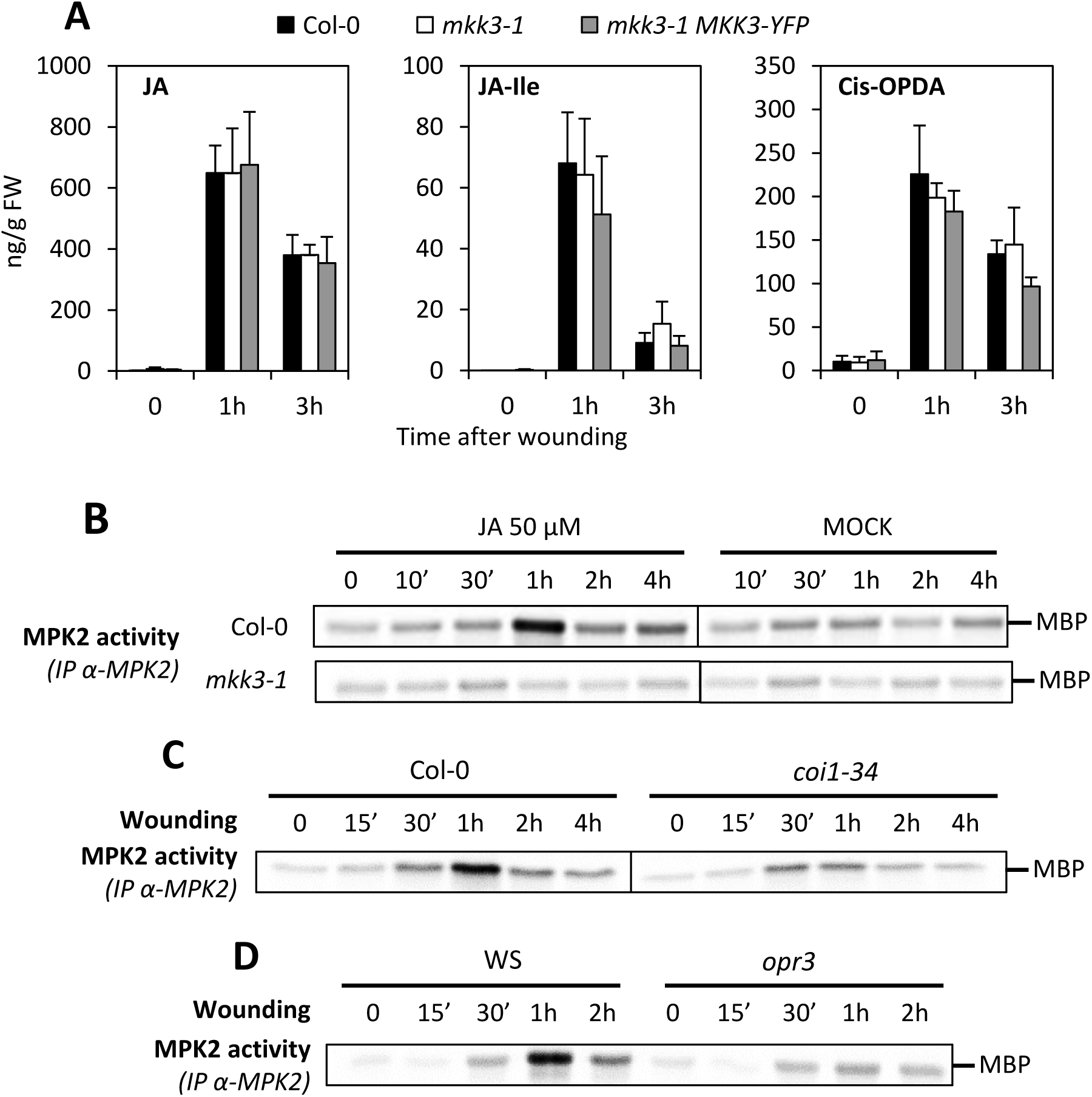
JA is involved in MKK3-MPK2 activation by wounding. **A.** JA, JA-Isoleucine and Cis-OPDA contents in wounded leaves of indicated genetic backgrounds. Values are mean±SE of 3 biological replicates. **B.** Kinase activity of MPK2 after immunoprecipitation with an anti-MPK2 specific antibody from Col-0 and *mkk3-1 in vitro* plantlets following 50µM JA or MOCK (EtOH) treatments. **C and D.** Kinase activity of MPK2 after immunoprecipitation with an anti-MPK2 specific antibody leaves of indicated genetic backgrounds following wounding at indicated times.

JA is sensed by CORONATINE INSENSITIVE1 (COI1) and *coi1* mutants are insensitive to JA (JA-Ile) (Katsir et al., 2008b, 2008a). To test whether the JA-induced activation of MPK2 is dependent on the COI1 receptor, we treated *coi1* mutant lines with JA and monitored subsequent MPK2 activity. JA-triggered MPK2 activation was reduced in the two mutant alleles, *coi1-34* and *coi1-16* (fig. S11). Moreover, in response to wounding, MPK2 activation was reduced in both *coi1-34* and *opr3*, a mutant impaired in JA biosynthesis (fig. 5C and 5D). Coherent with a role of JA in the signaling pathway, wound-triggered upregulation of clade-III *MAP3Ks* was also partially impaired in *coi1-34* (fig. S12). Taken together, these results indicate that JA synthesis and signaling, likely through the modulation of *MAP3K* expression, activate the MKK3-MPK2 module upon wounding.

### Rapid wound-induced activation of MPK3 and MPK6 is independent of MKK3

Classical stress-responsive MAPKs, such as Medicago SIMK and SAMK, tobacco SIPK and WIPK, and their Arabidposis homologs MPK3 and MPK6, are known to define functional modules which are activated by wounding (Meskiene et al., 2003; Seo et al., 2007; Li et al., 2017). In our conditions, an antibody raised against the ERK2 phospho-motif pT-E-pY (referred as anti-pTpY), detected two bands (around 40-45 KD) in Arabidopsis leaves rapidly after wounding (fig. 6A and S13A). In 15-minutes wounded leaves of *mpk6-2* and *mpk3-1* mutants, the higher and lower bands disappeared, respectively, confirming that MPK6 and MPK3 are activated by wounding (fig. S13B). Importantly, the wound-induced activation of MPK3 or MPK6 was not affected in *mkk3-1*, *mkk3-2*, *map3k14-1* or *map3k14-CR1* lines (fig 6A, S13D, S13E and S13F) but strongly impaired in the double mutant *mkk4mkk5* (fig. 6B, S13C) as previously described (Li et al., 2017). Moreover, MPK3 and MPK6 were not activated by JA and the wound-induced activation of MPK3/6 was not compromised in the JA-sensing-deficient *coi1-34* mutant (fig. S14). Overall, these data show that two MAPK modules are activated by wounding with different kinetics, a rapid MKK4/5-MPK3/6 module that is independent of JA, and a slower MKK3-MPK2 module that depends on JA signaling.

**Fig. 6.**
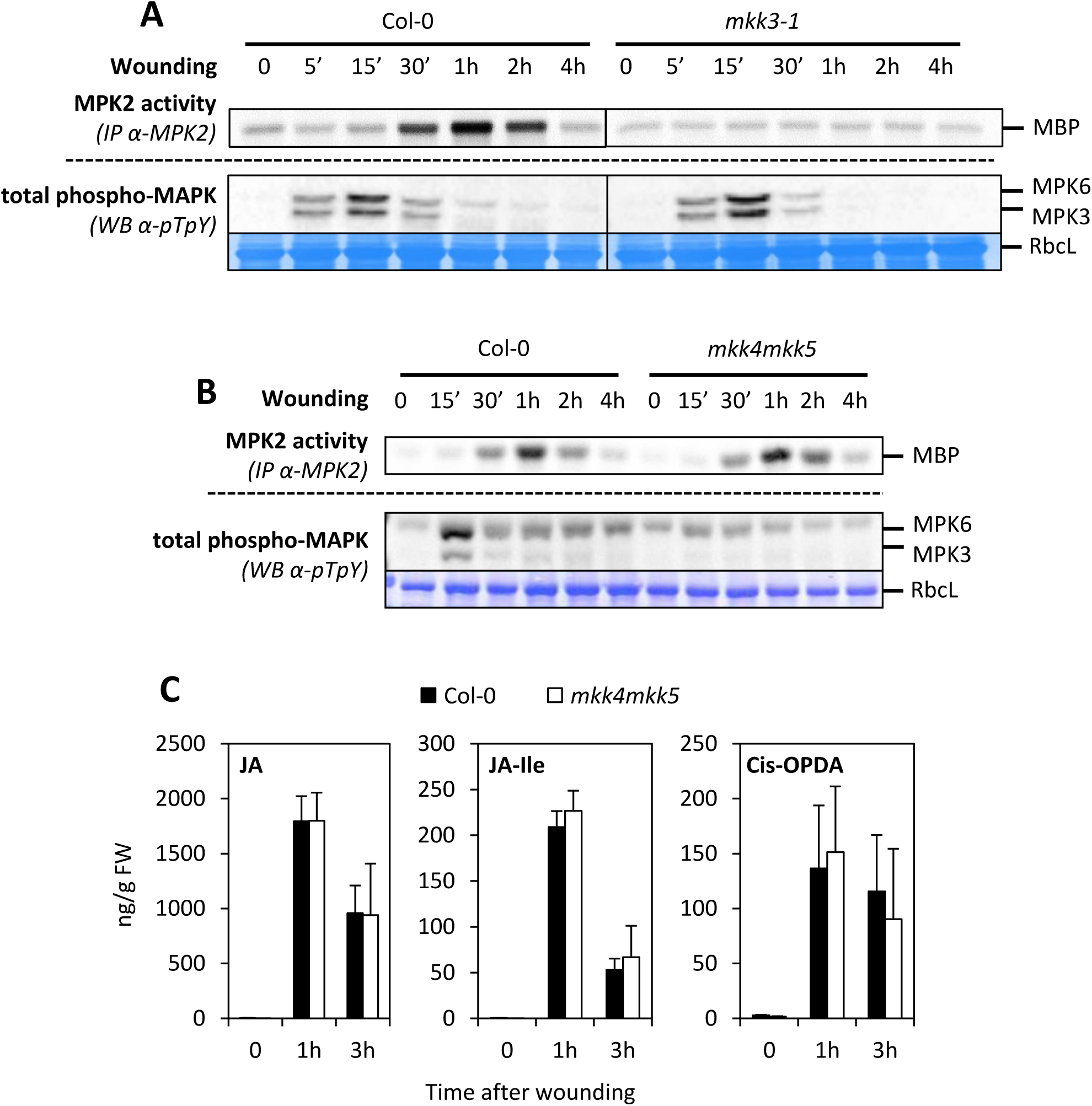
MKK4/5-MPK3/6 are not involved in wound-induced JA production. **A and B.** Kinase activity of MPK2 after immunoprecipitation with an anti-MPK2 specific antibody from Col-0 (A, B) and *mkk3-1* (A) and *mkk4mkk5* (B) leaves following wounding at indicated times. MPK3/6 activation was monitored by western-blot using antibody raised against the phosphorylated form of ERK2 (anti-pTpY). Equal loading is controlled by Coomassie staining. **C.** JA, JA-Isoleucine and Cis-OPDA contents in wounded leaves of Col-0 and *mkk4mkk5*. Values are mean±SE of 3 biological replicates.

### MKK3-MPK2 and MKK4/5-MPK3/6 modules are activated by wounding independently of each other

The fact that the MKK4/5-MPK3/6 module is activated rapidly might suggest that it could function upstream of MKK3-MPK2. This idea is supported by the finding that, in tobacco, SIPK and WIPK, which are homologues of *Arabidopsis* MPK3 and MPK6, were proposed to play a role in wound-induced JA synthesis (Seo et al., 1999; Heinrich et al., 2018; Seo et al., 1995, 2007) and we show here that JA is an important signal of wound-induced MKK3-MPK2 activation in Arabidopsis. To test whether the MKK4/5-MPK3/6 module acts upstream of MKK3-MPK2, MPK2 activation by wounding was compared in Col-0 and *mkk4mkk5* plants (fig. 6B). Although wound activation of MPK3 and MPK6 was barely detected in *mkk4mkk5* leaves, MPK2 activation was virtually unchanged. This result indicates that MPK2 activation does not rely on MKK4/5-MPK3/6 function. Although homologs of MPK3/6 in other species were proposed to control JA production, our results suggest that MPK3/6 are not involved in JA-mediated activation of MPK2 by wounding. To clarify this conundrum, we tested whether JA hormone synthesis was regulated identically in tobacco and Arabidopsis. For this purpose, we measured JA levels in Col-0 and *mkk4mkk5* plants at various time points after wounding (fig. 6C and fig. S15). JA, JA-Ile and cis-OPDA levels strongly and rapidly increased upon wounding but were unaffected in *mkk4mkk5* mutants, indicating that the MKK4/5-MPK3/6 module is not involved in wound-induced JA accumulation.

### The MKK3-MPK2 module is activated by Spodoptera littoralis feeding

Mechanical wounding partly mimics attacks by herbivorous insects. To test whether insects are able to activate the MAPK pathways with similar kinetics, starved *S. littoralis* larvae were allowed to feed on WT leaves for 15 minutes before removal. MAPK activation was subsequently monitored for 4h (fig. 7A). Under these conditions, MPK2 was activated after 30 minutes with a peak at 1 hour. MPK3 and MPK6 were only weakly activated at 5 and 15 min (fig. 7A). The MPK2 activation was dependent on MKK3 (fig. 7B) and COI1 (fig. 7C), like in wounding, suggesting that JA also plays a prominent role in the *S. littoralis*-triggered MPK2 activation.

**Fig. 7.**
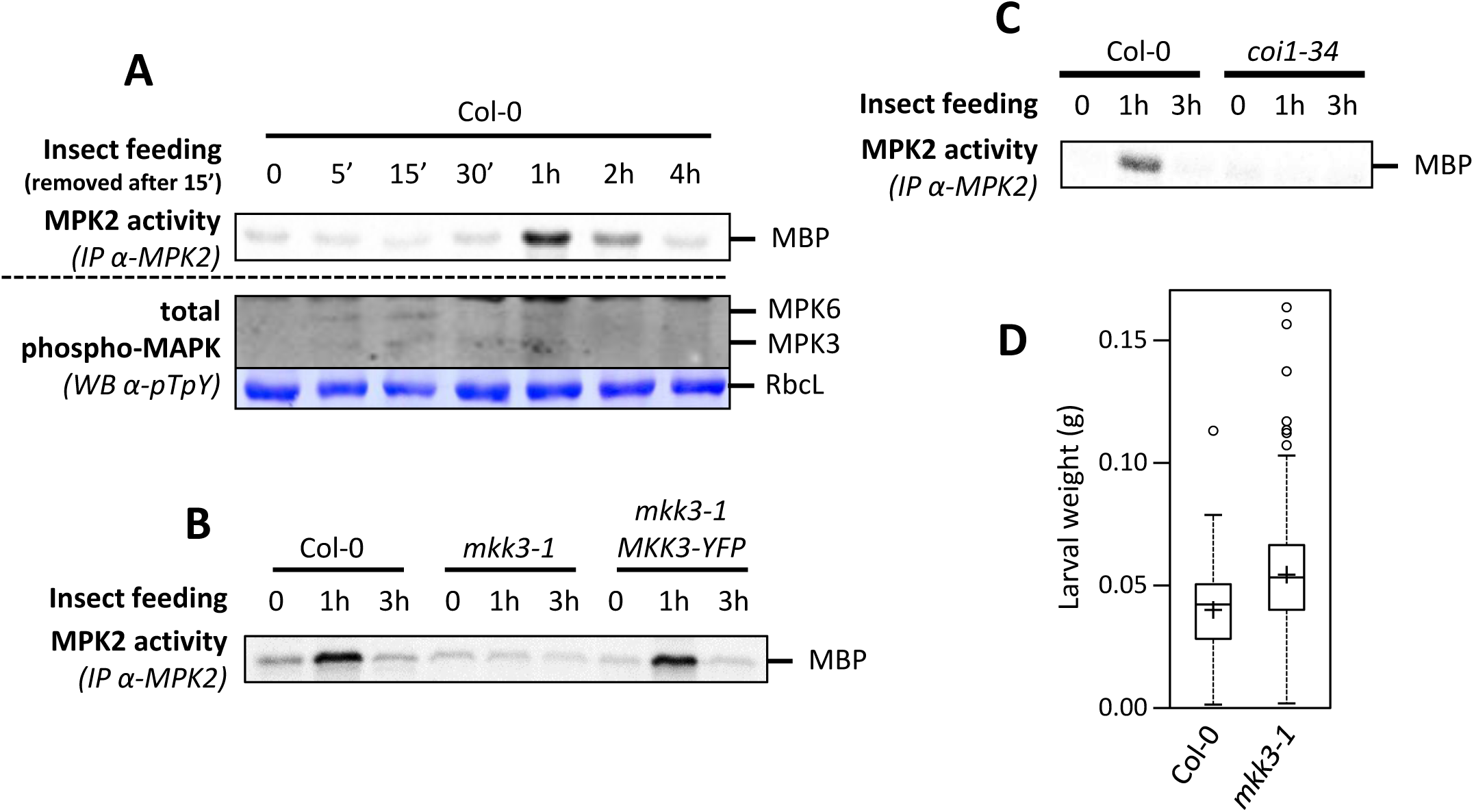
MKK3-MPK2 module is activated by *Spodoptera littoralis* feeding. **A.** Kinase activity of MPK2 after immunoprecipitation with an anti-MPK2 specific antibody from leaves on which *S. littoralis* fed during 15 minutes before to be removed (at t=15’). MPK3/6 activation was monitored by western-blot using antibody raised against the phosphorylated form of ERK2 (anti-pTpY). Equal loading is controlled by Coomassie staining. **B and C.** Kinase activity of MPK2 after immunoprecipitation with an anti-MPK2 specific antibody from leaves on which *S. littoralis* fed for 1 and 3 hours in Col-0 and *coi1-34*. **D.** Weight of *S. littoralis* caterpillars after feeding for 8 days on Col-0 and *mkk3-1* rosettes. Box plot shows distribution of caterpillar weight (n>120 in 5 biological replicates). Crosses show averages of biological replicates (39.5±5.2 and 54.4±6.6 mg for Col-0 and *mkk3-1*, respectively [n=5]; statistical difference based on the Mann-Whitney test with p<0.025).

To test whether the MKK3 module was necessary to restrict insect herbivory, we compared the growth of *S. littoralis* larvae on Col-0 and *mkk3-1* plants and observed a significant increase in larval weight when fed on *mkk3-1* leaves (fig. 7D). These results show that the MKK3 module can restrict *S. littoralis* growth.

## Discussion

We report in this study the activation of two independent MAPK modules by wounding in Arabidopsis plants (fig. 8). The first module is defined by MKK4/5-MPK3/6. The second module, defined by MKK3-clade-C MAPKs, was suspected from previous preliminary information (Ortiz-Masia et al., 2007). These two modules are activated independently of each other with very distinct kinetics. We also demonstrated that jasmonic acid, which is produced in response to wounding and herbivores, is an important mediator of the activation of the MKK3 module. Overall, this work provides insight into wound signaling of Arabidopsis.

**Fig. 8.**
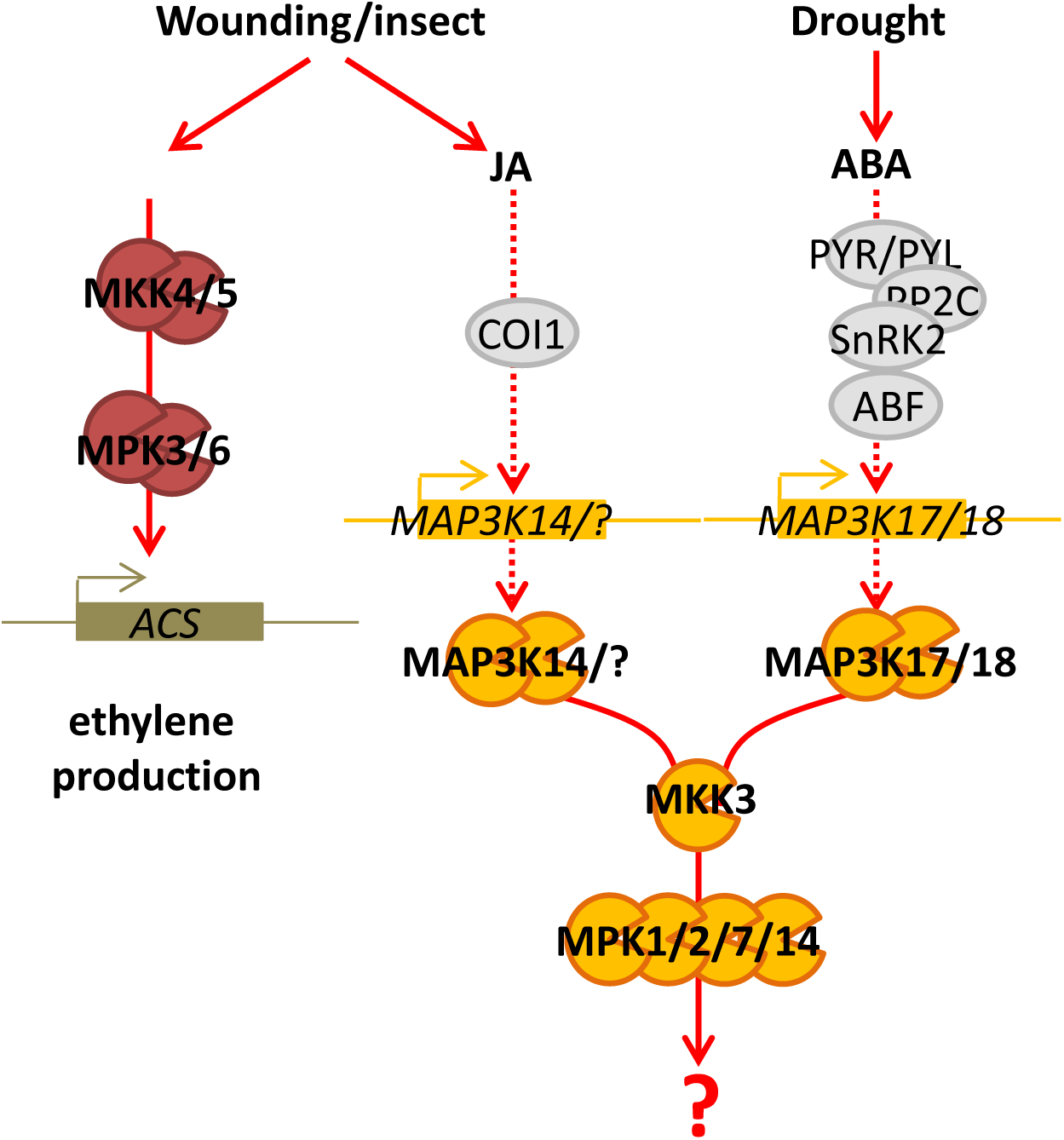
Working model of MAPK activation by wounding. Wounding and insect feeding activate two MAPK modules: a rapid one composed of MKK4/5-MPK3/6 which regulates notably ethylene production and a slow one composed of clade-III MAP3Ks-MKK3-MPK1/2/7/14 whose activation is under the control of a JA-dependent production of MAP3Ks. Drought also activates MAP3Ks-MKK3-MPK1/2/7/14 through an ABA-dependent step.

### MAP3K13/14/15/16/17/18/19/20-MKK3-MPK1/2/7/14: general late responsive MAPK modules?

Our data suggest that the MKK3-MPK2 module is activated through the JA- and wound-induced transcription of *s*everal clade-III MAP3Ks. Among them, *MAP3K14* shows the highest and earliest expression upon wounding. Coherently, we observed a reduction of MPK2 activation by wounding specifically at early time points in the loss-of-function *map3k14* CRISPR lines. We also observed a stronger wound-induced MPK2 activation in *map3k14-1*, which expresses a truncated and more stable form of MAP3K14. In addition, other members of the clade-III MAP3Ks, such as *MAP3K18*, *MAP3K17*, *MAP3K19* and *MAP3K20*, which are induced by wounding at a later time point, may maintain MPK2 activity after MAP3K14-triggered initial activation. Among the 8 clade-III MAP3K proteins, only MAP3K13 and MAP3K14 have a carboxyl-terminal extension which could correspond to a transmembrane domain, possibly suggesting a particular role at the membrane, a location thought to play a role in cellular wound sensing (Farmer et al., 2014). Unfortunately, we did not succeed to see any fluorescent signal in lines expressing MAP3K14::YFP after wounding, suggesting a very low expression level of MAP3K14 or alternatively that the YFP fusion at the C-terminus may reduce MAP3K14 stability (data not shown). As expected in a process requiring *de novo* protein production, the general inhibitor of protein synthesis, cycloheximide, fully abolished the wound-induced MPK2 activation. In the past, de novo synthesis of MAP3Ks has been well documented in the case of the MKK3-MPK7 activation by drought which requires ABA-dependent MAP3K18 protein accumulation (Danquah et al., 2015; Boudsocq et al., 2015). Here we describe another stress triggering MKK3 module activation by *de novo* protein synthesis of MAP3Ks via transcriptional regulation. This new result comforts this model, suggesting that the late responsive signaling MAPK pathways are more general than first thought. Therefore the module could broadly regulate a second layer of events in response to environmental constraints (Colcombet et al., 2016).

MKK3 modules have been by far less characterized than the iconic stress-activated module MKK4/5-MPK3/6. An obvious explanation lies in its rather atypical slow activation, which was unexpected; textbooks often teach that MAPK modules are early/rapid signaling actors. Additionally, clade-C MAPKs have never been detected so far by *in-gel* kinase assays. Nevertheless, phosphorylated activation motifs of clade-C MAPKs have been found in phosphoproteomic approaches suggesting that they are often activated. For example, MPK1/2 phosphorylation was reported in response to ABA (Umezawa et al., 2013) and DNA damage-inducing irradiations (Roitinger et al., 2015). Interestingly, in *map3k14-1* that over-activates MPK2 in response to wounding, the anti-pT-E-pY antibody detected a new band appearing below MPK3 and MPK6 with a kinetics fitting clade-C MAPK activation (Fig. S13E). This suggests that clade-C MAPKs are tightly controlled to avoid full activation. More exhaustively, transcriptional regulation of clade-III MAP3Ks by stresses seems to occur very often based on expression databases (Winter et al., 2007; Zimmermann et al., 2004), and suggests important roles of MKK3 modules in environmental perception. Notably, MAP3K13, MAP3K18 and MAP3K20 are induced by a large number of environmental signals including osmotic, salt and drought stresses. Whereas MKK3 is a hub of the module, the related MAP3Ks and MAPKs are encoded by multigenic families. The input signal specificity is conferred by the transcriptional regulation of the *MAP3K* genes but we cannot exclude that MAP3K activity is also directly modulated by input stresses, as suggested in ABA signalling (Matsuoka et al., 2015; Mitula et al., 2015). The situation is less clear for MAPKs: MPK1/2/7/14 seem to be activated identically by a stress, based on protoplast experiments and immunoprecipitation from organs (Danquah et al., 2015). Our hypothesis to explain this functional redundancy is that they may be expressed in different cells/compartments or target specific substrates. This should be an important point to address in the future.

Last, MKK3 has also been proposed to function upstream of MPK6 and MPK8 in various contexts such as blue light, dark-light transition and ROS homeostasis (Sethi et al., 2014; Takahashi et al., 2007, 2011; Lee, 2015). We previously reported that these functional connections were not found when combining these kinases in protoplast expression systems (Danquah et al., 2015). Additionally, in the present work, we did not see any MKK3-MPK6 functional connection in the context of wounding. Nonetheless, it is possible that such connections exist in other physiological contexts in specific cell types or organs (Colcombet et al., 2016).

### Is wound-induced JA production connected to MAPK modules?

The phytohormone JA plays a critical role in the wound response (Wasternack, 2018). It is massively produced in a few minutes after wounding by chloroplast- and perixisome-localized biosynthetic enzymes and is involved both in local and long-distance responses (Wasternack and Hause, 2013; Koo et al., 2009). Plants impaired in JA synthesis and signaling have weakened responses to wounding and are much more susceptible to chewing insects (Howe and Jander, 2008). The JA core signaling module is well described (Thines et al., 2007; Chini et al., 2007): JASMONATE ZIM DOMAIN (JAZ) proteins sequester transcription factors, notably basic helix-loop-helix (bHLH) factors such as MYC2 to prevent them from activating JA-responsive genes. The binding of JA-Ile to COI1 receptor, which is an F-Box E3 ligase, increases COI1 affinity for JAZ proteins, triggering their ubiquitination and consequent proteasome-dependent degradation. The released transcription factors become free to modulate gene expression (Thines et al., 2007). In this article, we report a reduction of wound-triggered MPK2 activation in *opr3* and *coi1* mutants, which are impaired in JA synthesis and signaling, respectively. Therefore we can conclude that the activation of the MKK3-MPK1/2/7/14 module is downstream of JA signaling and synthesis. The remaining activation in these mutants could be explained either by the existence of JA-independent pathways controlling MAP3K transcriptional regulation in response to wounding or by the fact that they are not total loss-of-function mutants. *opr3* mutant was reported to still produce some JA-Ile (Chini et al., 2018) and *coi1-34* is not a full loss-of-function allele (Acosta et al., 2013). Surprisingly, the wound-triggered up-regulation of *MAP3K* transcripts is only mildly impaired in the JA mutants whereas the MAPK activation is strongly reduced. It is possible that the active MAP3K amount is limiting, titrated by competition. Alternatively, beside transcriptional induction, MAP3Ks may also require a JA-dependent modification (i.e. phosphorylation) for their activation, and the two impaired regulations would lead to a strong reduction of MPK2 activation. This last hypothesis is supported by the fact that MAP3K18 intrinsic activity increases upon ABA treatment (Matsuoka et al., 2015; Mitula et al., 2015).

These results add a new layer of complexity in the chain of events triggered by JA perception. The question of what is controlled by the JA-activated MKK3 module remains open. MKK3-related modules had been shown to regulate ROS homeostasis in response to wounding, but was then proposed to work upstream of the clade-D MAPK MPK8 (Takahashi et al., 2011). Recently, the MAP2K inhibitor PD98059 was shown to reduce JA-triggered activation of enzymes involved in ascorbate and glutathione metabolism in maize leaves (Shan and Sun, 2018). Overall, these results point to redox homeostasis as a putative target of MKK3 module in the general response to wounding and JA.

The signaling pathway from wound signaling to JA production is largely unknown. One known event is the transient increase of cytosolic Ca^2+^ (Maffei et al., 2004; Kiep et al., 2015) which must be decoded by Ca^2+^-sensing proteins. Two calmodulin-like-proteins, CML37 and CML42, have been identified as positive and negative regulators, respectively, of JA-mediated defense in Arabidopsis plants after herbivore attack (Vadassery et al., 2012; Scholz et al., 2014). While CML42 very likely affects the binding of JA-Ile to the receptor, CML37 regulates the expression of *JAR1*, the enzyme catalyzing the JA-Ile formation. Our work shows that JA synthesis is not under the control of MPK3/6. This is surprising as the homologs of MPK3 and MPK6, WIPK and SIPK, respectively, were shown to modulate JA levels in response to wounding in tobacco (Seo et al., 1999, 1995, 2007; Heinrich et al., 2018). This apparent discrepancy might be explained by different approaches used in the studies. Notably, we took advantage of a unique *mkk4mkk5* mutant issued from a tilling screen (Zhao et al., 2014), in which the wound-induced activation of MPK3/6 is strongly reduced but not fully abolished. It is possible that the remaining weak activation of MPK3/6 is sufficient to trigger downstream events such as the hormonal production. Nevertheless, this hypothesis is unlikely as this mutant is clearly impaired in wound-induced ethylene production (Li et al., 2017). Alternatively, and more interestingly, these results suggest that homologous MAPKs may not regulate the same responses in various plant species, despite being activated similarly by a given stress. This might correspond to diverging adaptation strategies in tobacco and Arabidopsis.

MPK3 and MPK6 are very well-known stress-responsive MAPKs. Many of their substrates have been identified in the context of Microbe-Associated Molecular Pattern (MAMP) signaling (Bigeard et al., 2015; Rayapuram et al., 2018; Bigeard and Hirt, 2018). However, very little is known about their functions in response to other stresses and notably in response to wounding. The recent discovery that the MKK4/5-MPK3/6 module is involved in wound-induced ethylene production through the transcriptional regulation of *ACS* genes was the first report of their role in wound signaling in Arabidopsis (Li et al., 2017). Interestingly, the same module is involved in ethylene production in response to flg22 and Botrytis (Li et al., 2012; Liu and Zhang, 2004). This regulation occurs through both a direct phosphorylation-dependent stabilization of the ACS6 enzyme and the up-regulation of *ACS* genes by the phosphorylation-dependent activation of the WRKY33 transcription factor. In rice, OsMPK1, the closest homolog of AtMPK6, was also shown to interact with and phosphorylate WRKYs *in vitro* (Yoo et al., 2014). Interestingly, cycloheximide, which is a potent blocker of MKK3 module activation (fig. 3), is a strong activator of MPK3/6 (fig. S16) and triggers the transcriptional up-regulation of a large set of MAMP-regulated genes (Navarro et al., 2004). Overall, these data suggest that the MKK4/5-MPK3/6 module may be involved in the control of gene expression in response to wounding as for other stresses (Frei dit Frey et al., 2014).

### Roles of the MAPKs in the interaction with herbivores

MAPKs also play an important role in plant-herbivore interactions (Hettenhausen et al., 2017). Early studies in *Nicotiana tabaccum* showed that WIPK is activated rapidly by wounding and that WIPK-silenced plants have reduced levels in defense-related genes and JA (Seo et al., 1995). In *Nicotiana attenuata*, mechanical wounding and oral secretion of the herbivore *Manduca sexta* activate SIPK and WIPK within 5 minutes (Wu et al., 2007). In Arabidopsis, grasshopper oral secretion increases the wound-induced activation of MPK3 and MPK6 (Schäfer et al., 2011). In *Solanum lycopersicum, M. sexta* feeding also activates WIPK and SIPK homologs SlMPK3 and SlMPK1/2, respectively, and silencing SlMPK1/2 reduces herbivory-induced JA levels resulting in enhanced larval growth (Kandoth et al., 2007). However, other studies in *N. tabaccum* and *N. attenuata* reveals more complexity in the JA-MAPKs relationship depending on herbivores. In response to wounding, tobacco MPK4 is activated in minutes and silencing NtMPK4 compromises JA-responsive gene induction (Gomi et al., 2005). By contrast, silencing NaMPK4 does not affect JA production nor resistance to the generalist herbivore *Spodoptera littoralis*, but increases resistance to *M. sexta* (Hettenhausen et al., 2017). This indicates that some functions of MPK4 are specific to particular herbivore interactions and that further studies will be necessary to clarify the role of different MAPK modules in wound- and herbivory-induced JA signaling.

## Methods

Primer sequences for plant genotyping, molecular cloning and expression analysis are provided in the supplemental table 1.

### Plant material

*mkk3-1, coi1-34* (Acosta et al., 2013), *opr3*, *mpk3-1*, *mpk6-2* (Galletti et al., 2011), *mkk4mkk5* (Su et al., 2017) and *map3k17map3k18:MAP3K18locus-YFP* (Danquah et al., 2015) were published previously. *mkk3-2* (Salk_208528) and *map3k14-1* (Gabi_653B01) were identified in public databases in the frame of this work.

To generate CRIPSR/Cas9 mutants of *MAP3K14* gene, CRISPR guide sgRNA targeting MAP3K14 was generated by cloning Cs9-3K14-F/Cs9-3K14-R primers into pDGE65 according to Ordon et al. (2016). The vector was transformed into *Agrobacterium tumefaciens* C58C1 and used to transform Arabidopsis Col-0 plants using floral dipping (Clough and Bent, 1998). The number of inserted T-DNA in 20 independent Basta-resistant lines were identified by segregation of T1 seeds *in vitro*. To eliminate the Cas9 endonuclease, the Basta-sensitive plantlets were rescued by careful transfer on pots and, after recovery, genotyped for mutations in *MAP3K14*, for the absence of mutations in the closest homologue gene *MAP3K13* and for the absence of the pDGE65 T-DNA in the genome. Two independent and homozygous lines (*MAP3K14–CR1* and *MAP3K14–CR2*) were identified. MPK1, MPK2, MPK7 and MKK3 loci upstream of the STOP codon and containing 5’UTR, introns and exons as well as a large part of promoters were amplified using appropriate primers (Table S1) and the iProof enzyme (Bio-Rad), digested with appropriate restriction enzymes and fused in frame with HA or YFP tags in home-made binary vectors derived from pGREEN0229, referred as pGREEN0229-MPK1-HA, pGREEN0229-MPK2-HA, pGREEN0229-MPK7-HA and pGREEN0229-MKK3-YFP. Vector sequences are provided as table S2. Vectors were transformed into *A. tumefaciens* strain C58C1 containing pSOUP helper plasmid (Hellens et al., 2000). Kanamycin-resistant Agrobacteria were then used to transform *Col-0* or *mkk3-1* Arabidopsis plants using the floral dipping method (Clough and Bent, 1998). Segregation analysis of Basta-resistance was used to identify homozygous lines.

### Wounding experiments

A detailed protocol for wounding is provided (text S1). Briefly, single plants were grown on pellets (Jiffy-7 38mm Pellet-Pack, Ref # 32204011, Jiffy Products International AS, Norway) in a Percival growth chamber with 12 hours dark/12 hours light at 22°C and 70% humidity for 4 weeks before experiments. Three fully-expended leaves of an Arabidopsis rosette were wounded three times using tweezers (5 mm round tip with line serrated surfaces). Wounding was performed at proper time to obtain the expected incubation duration with the harvesting of samples (3 leaves × 3 plants) at 3-4 pm. For cycloheximide treatments, a solution containing 100 µM CHX (Sigma-Aldrich, Ref# C4859) in 0.03% DMSO as well as a mock (0.03% DMSO) were sprayed equally onto rosettes 150 min before collecting samples. Samples were frozen in liquid nitrogen, grinded either using pestles and mortars or homogenizer (SIGMA) and kept at −80°C before further investigation. Experiments were typically repeated 3 times with plants grown independently.

### JA experiments

For JA-treated samples, seeds were sterilized (15 min in 70% EtOH) and 20-30 plantlets were grown in small petri dishes (diameter 6 cm) containing 8 mL ½ MS + 1% sucrose for 10 days with 16 h light/8 h night at 22°C. Then plantlets were treated with 0.1% ethanol (mock) or 50 µM JA (Sigma-Aldrich, ref # J2500). To stop treatments, plantlets were rapidly dried and frozen in liquid nitrogen.

### Insect experiments

Larvae of the generalist lepidopteran species *Spodoptera littoralis* were reared as previously published (Vadassery et al., 2012). To monitor MPK2 activation or measure oxylipin production in response to insect, 4^th^ instar larvae were starved overnight prior to plant feeding for 1h and 3h. To monitor MPK3/MPK6 activation, larvae were removed after 15 minutes. For long-term feeding assays (8 days), first instar larvae were used according to Vadassery et al. (2012). When different plant lines were used at the same time, they were kept separated from each other to avoid any kind of contact and placed randomly in the experimental setup.

### Kinase Assays

A detailed kinase assay protocol is provided (text S1). Kinase assays using α-MPK2 (Ortiz-Masia et al., 2007), α-HA (SIGMA A2095), α-MPK3 (SIGMA M8318) and α-MPK6 (SIGMA A7104) antibodies as well as western-blots were performed as previously described (Danquah et al., 2015; Ortiz-Masia et al., 2007). It should be noted that α-MPK2 antibodies can specifically immunoprecipitate MPK2 although they fail to detect the protein on western-blots of total proteins (Ortiz-Masia et al., 2007).

### Other assays

For gene expression analysis, plants were collected at indicated times and frozen in liquid nitrogen. RNA extraction, cDNA synthesis and qRT-PCR analysis were performed as previously described using primers in table S1 (Danquah et al., 2015). Yeast 2-hybrid assays were performed as previously described except that the selective medium was generated with 0.2% dropout-T-L-U [US Biological] (Berriri et al., 2012). Jasmonate contents in *Arabidopsis thaliana* leaves were analyzed by LC-MS as described previously (Almeida-Trapp et al., 2014).

### Accession Numbers

Sequence data from this article can be found in the Arabidopsis Genome Initiative or GenBank/EMBL databases under the following accession numbers: At1g10210 (MPK1), At1g59580 (MPK2), At3g45640 (MPK3), At2g43790 (MPK6), At2g18170 (MPK7), At5g40440 (MKK3), At1g51660 (MKK4), At3g21220 (MKK5), At1g07150 (MAP3K13), At2g30040 (MAP3K14), At5g55090 (MAP3K15), At4g26890 (MAP3K16), At2g32510 (MAP3K17), At1g05100 (MAP3K18), At5g67080 (MAP3K19) and At3g50310 (MAP3K20).

## Author Contributions and Acknowledgments

C.S., H.H., and J.C. designed the research. C.S., S.T.S., M.B., C.C., A.K., M.A-T., A.M. and J.C. performed research. All the authors contributed to the manuscript.

We thank the Stress Signaling group for critical discussion of this work. We also thank Shuqun Zhang and Edward Farmer for providing *mkk4mkk5* and *coi1-34* seeds. This work has benefited from a French State grant (LabEx Saclay Plant Sciences-SPS, ANR-10-LABX-0040-SPS), managed by the French National Research Agency under an “Investments for the Future” program (ANR-11-IDEX-0003-02). C.S. and C.C. were funded by SPS PhD fellowships. Marilia Almeida-Trapp gratefully acknowledges financial support by a Capes-Humboldt Research Fellowship. This publication has been written with the support of the AgreenSkills+ fellowship program which has received funding from the EU’s Seventh Framework Program under grant agreement No. FP7-609398 (AgreenSkills+ contract) to STS.

## Figure legends

**Fig. S1.**
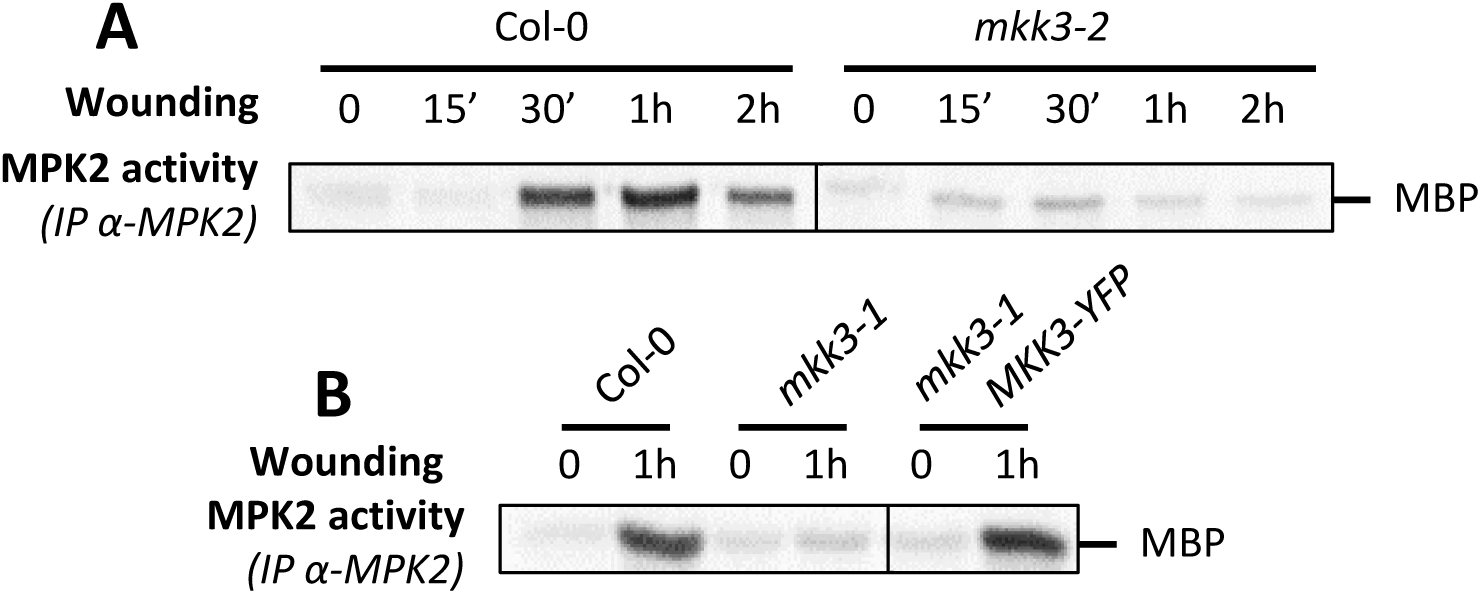
MPK2 activation by wounding depends on MKK3. **A and B.** Kinase activity of MPK2 after immunoprecipitation with an anti-MPK2 specific antibody from leaves of Col-0 and *mkk3-2* KO (A) and Col-0, *mkk3-1* KO and *mkk3-1 MKK3-YFP* complemented line (B) following wounding at the indicated times.

**Fig. S2.**
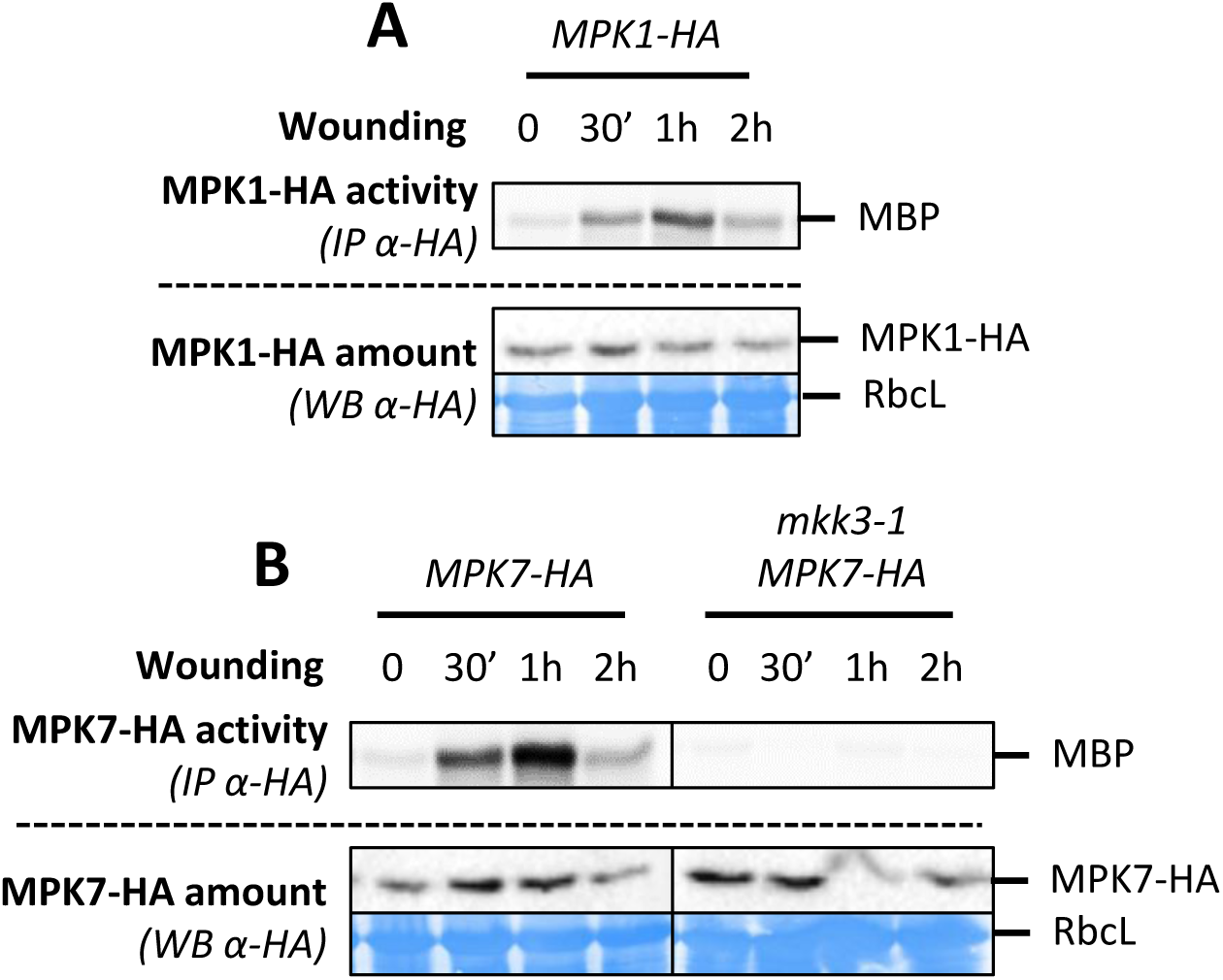
MPK1 and MPK7 are activated by wounding. **A and B.** Kinase activity of MPK1 (A) and MPK7 (B) after immunoprecipitation with an anti-HA antibody from leaves of plants expressing a HA-tagged version of MPK1/7 following wounding at the indicated times. Protein amount is monitored by western-blot using anti-HA antibody. Equal loading is controlled by Coomassie staining.

**Fig. S3.**
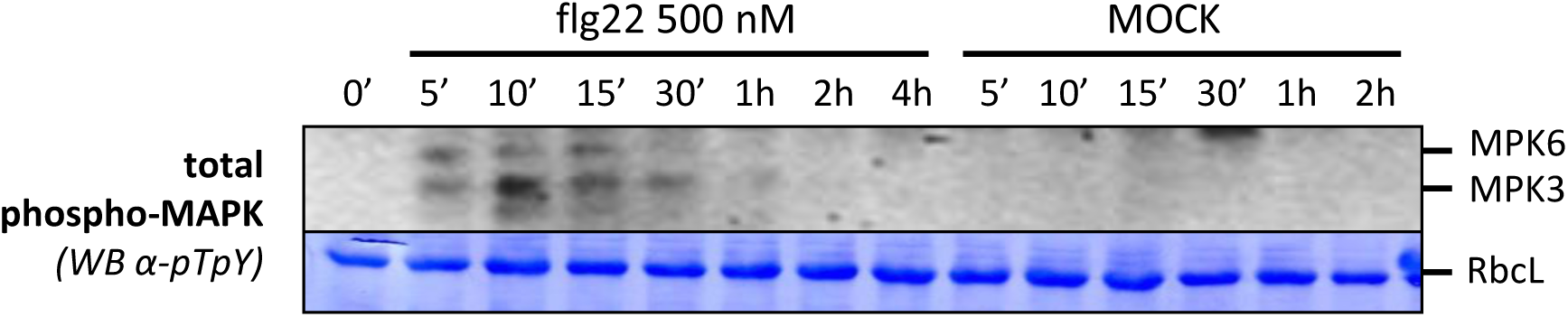
Flg22 rapidly and transiently activates MPK3 and MPK6 in leaf punches. Punches from Col-0 leaves were equilibrated over night in water and then treated for the indicated time with 500 nM flg22 or water (MOCK). MPK3 and MPK6 phosphorylation is monitored by western-blot using an antibody raised against the phosphorylated form of ERK2 (anti-pTpY). Equal loading is controlled by Coomassie staining.

**Fig. S4.**
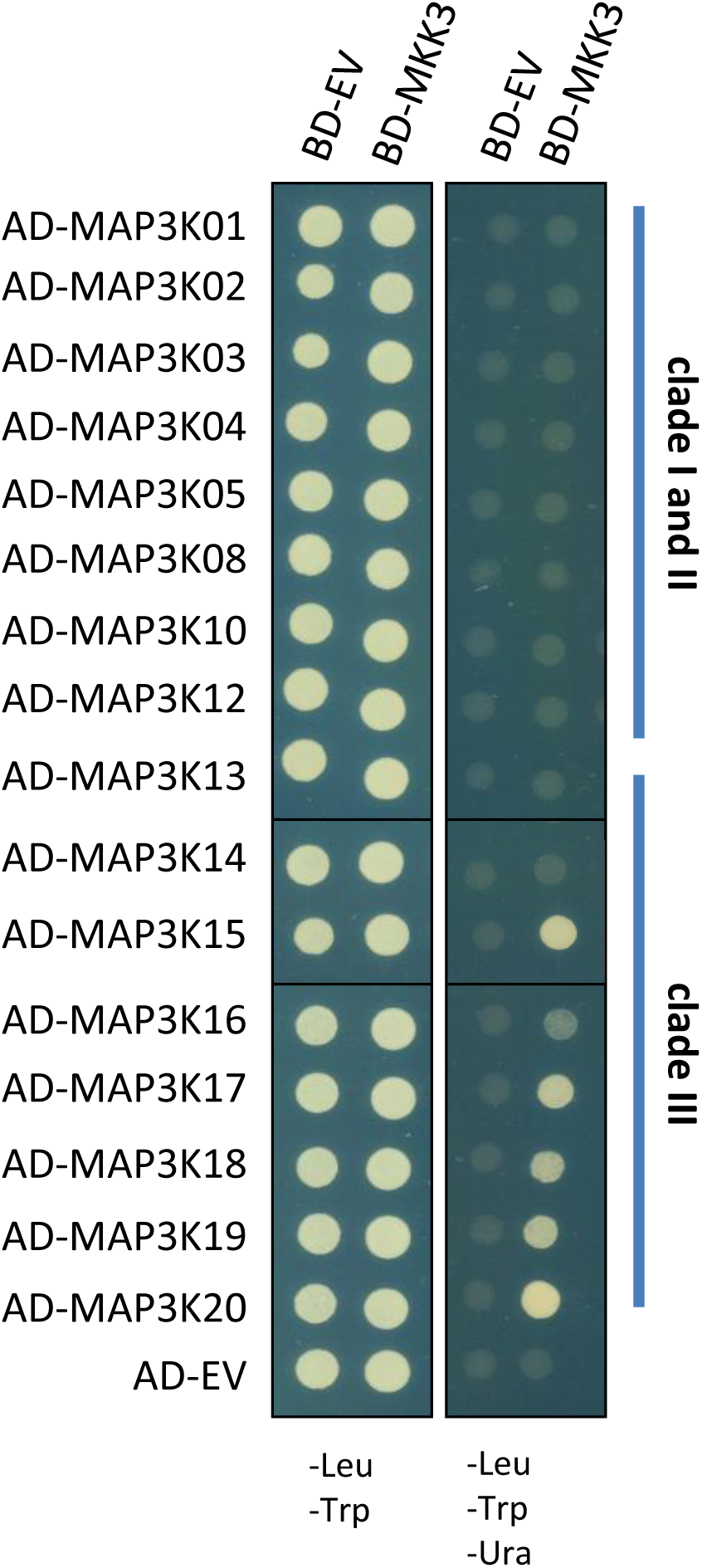
MKK3 and clade-III MAP3Ks interact in yeast 2-hybrid assay. Yeast 2-hybrid experiment showing the interaction of MEKK-like MAPKs with MKK3. 16 of the 20 MEKK-like kinases of *Arabidopsis* and MKK3 were respectively fused to the activation domain (AD) and the DNA binding domain (BD) of a transcription factor allowing the growth of yeast on a selective medium. EV stands for Empty Vector.

**Fig. S5.**
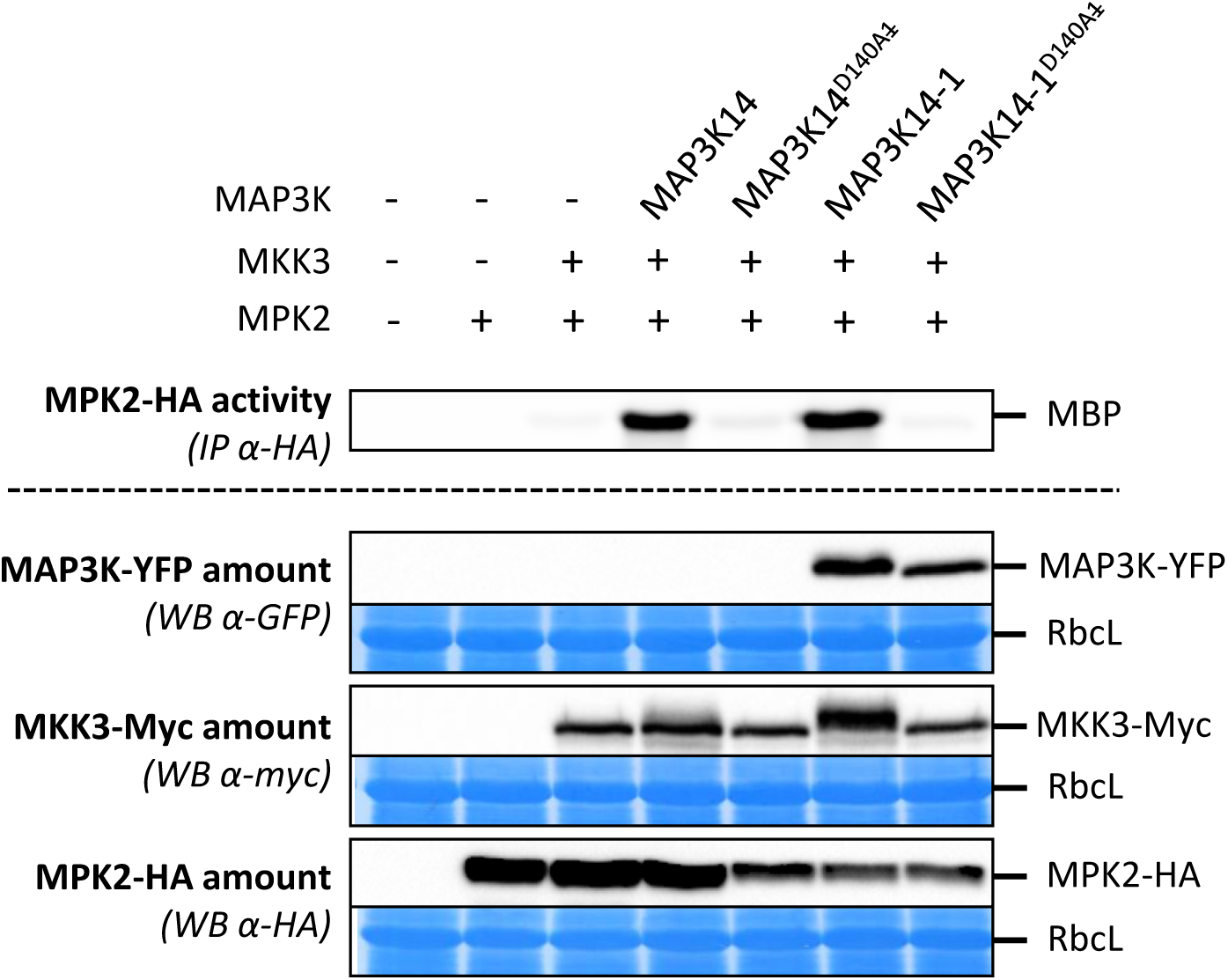
MAP3K14 activity is necessary for MKK3-MPK2 activation. Kinase activity of HA-immunoprecipitated MPK2 expressed in *mkk3-1* mesophyll protoplasts in the presence or absence of MKK3 and indicated versions of MAP3K14. MAP3K14-1 is a truncated and more stable form of MAP3K14 (see fig. S8). The detection of MAP3K14-1 and MAP3K14-1^D140A^ by western-blot strengthens the results obtained with MAP3K14. Western-blots show protein expression levels. Equal loading is controlled by Coomassie staining.

**Fig. S6.**
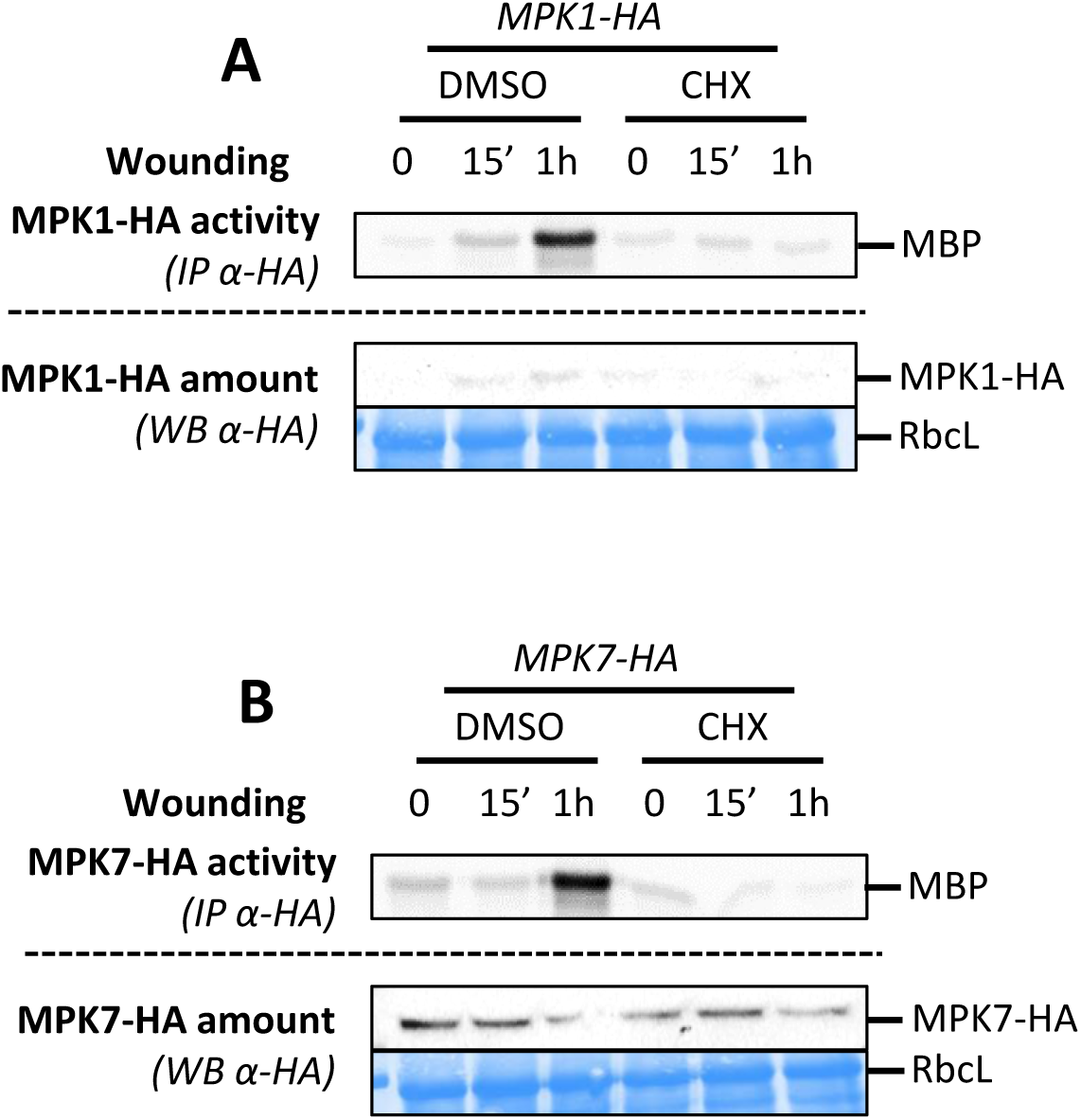
MPK1 and MPK7 activations by wounding require protein synthesis. **A and B.** Kinase activity of MPK1 (A) and MPK7 (B) after immunoprecipitation with an anti-HA antibody from leaves of indicated genetic background following 100µM CHX and MOCK (DMSO) spraying prior to wounding. Protein amount is monitored by western-blot using an anti-HA antibody. Equal loading is controlled by Coomassie staining.

**Fig. S7.**
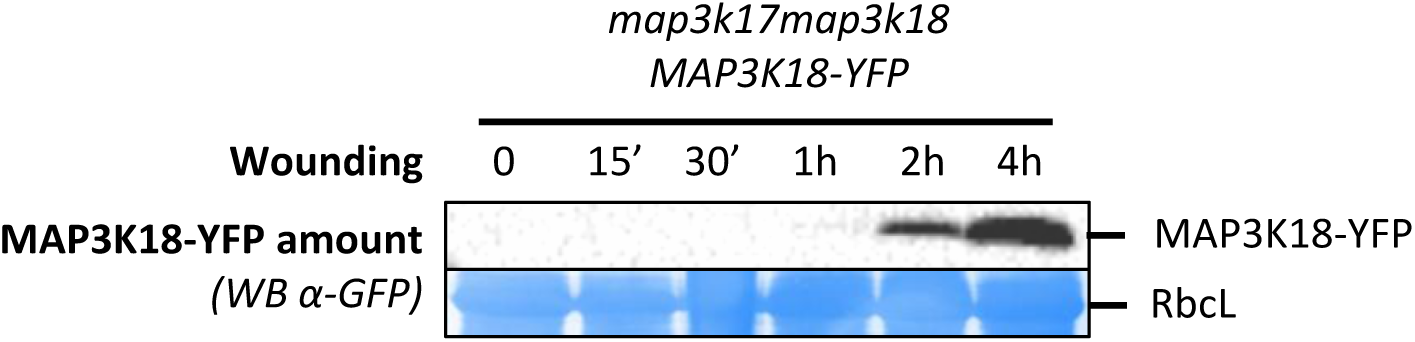
MAP3K18 accumulates in response to wounding. Western-blot using anti-GFP antibody showing MAP3K18-YFP protein level expressed under its own promoter in wounded leaves. Equal loading is controlled by Coomassie staining.

**Fig. S8.**
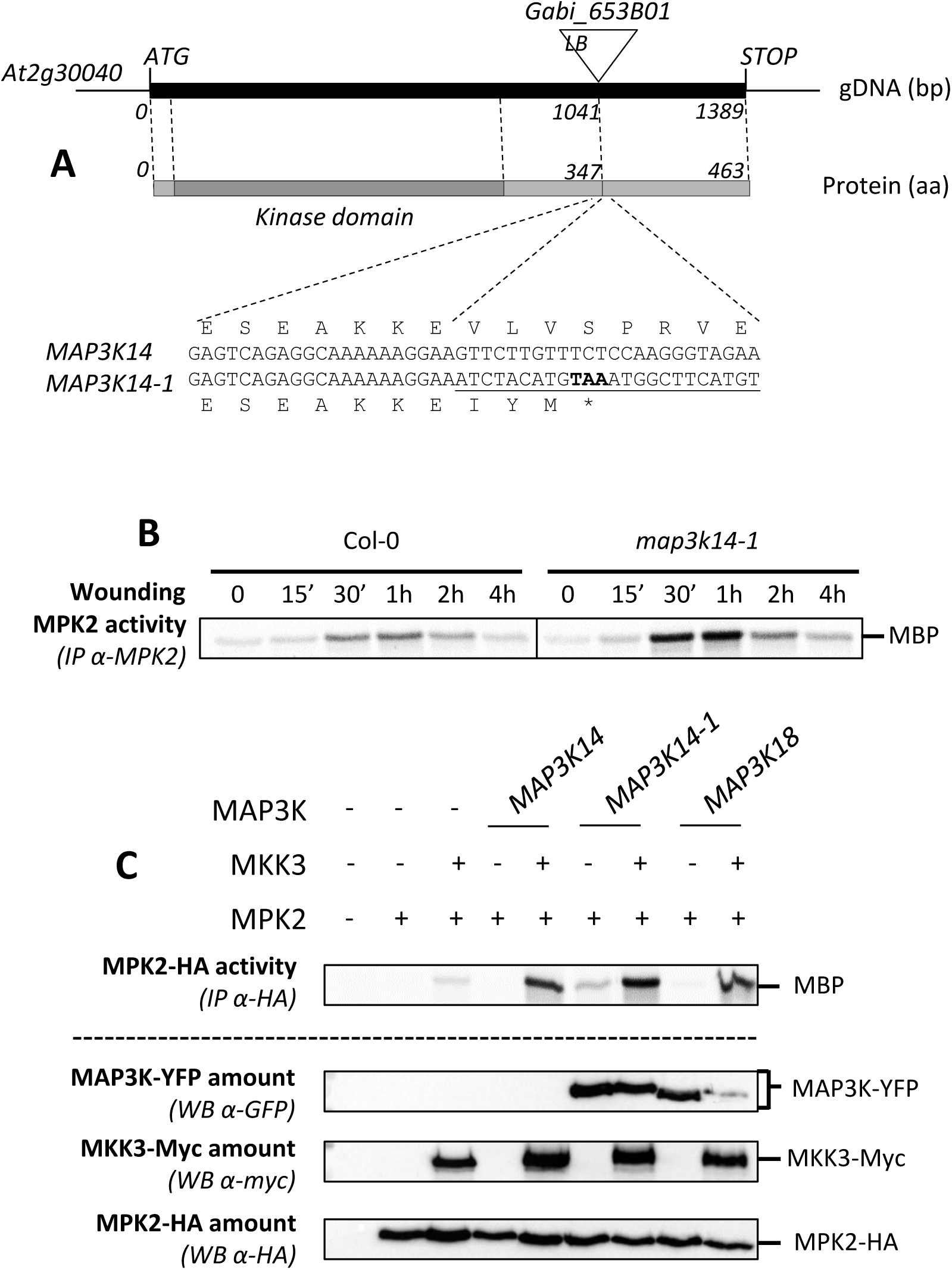
MPK2 Activation by wounding is higher in *map3k14-1*. **A.** Genomic structure of T-DNA insertion in *map3k14-1*. The resulting protein MAP3K14-1 is shorter of 116 amino acids. **B.** Kinase activity of MPK2 after immunoprecipitation with an anti-MPK2 antibody from WT and *map3k14-1* leaves following wounding. **C.** Kinase activity of HA-immunoprecipitated MPK2 expressed in *mkk3-1* mesophyll protoplasts in the presence or absence of MKK3 and MAP3K14wt and MAP3K14-1. Western-blots show protein expression levels.

**Fig. S9.**
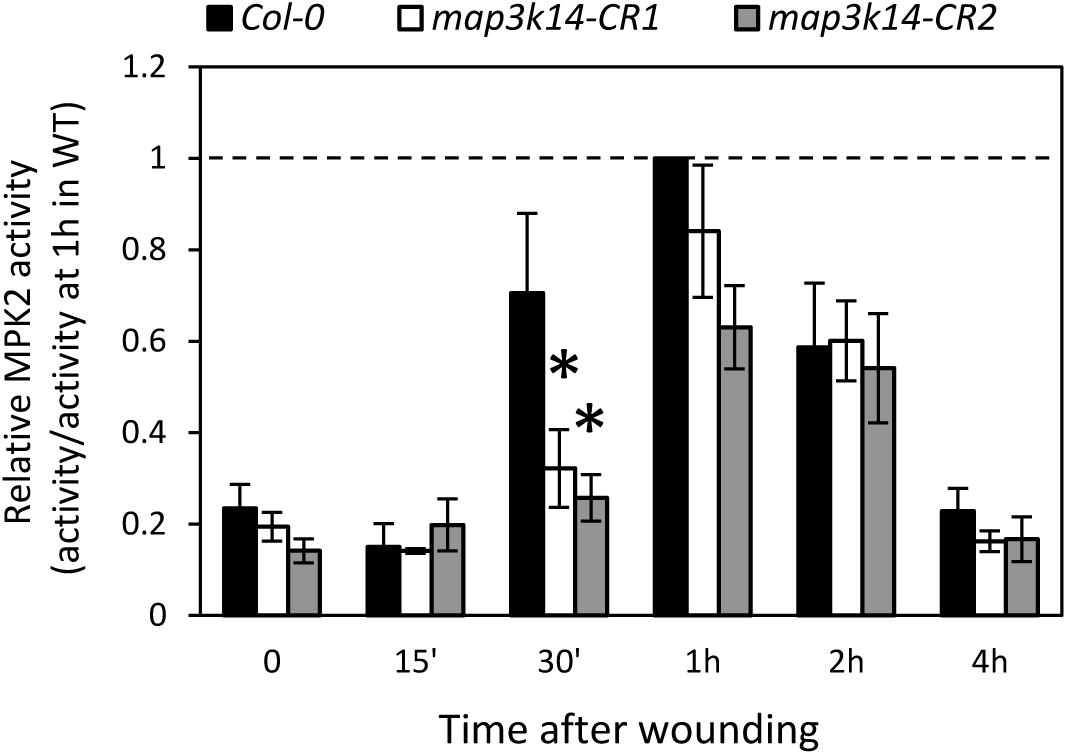
Quantification of MPK2 activation by wounding in *map3k14-CR1* and *-CR2* lines. Quantification of MPK2 activity in response to wound in Col-0 and 2 independent *map3k14-CR* lines. For each experiment (such as in fig 4B), phosphorylation values were normalized to the phosphorylation value at 1 hr in WT. Values are mean±SE of 4 biological replicates. Stars indicate statistical differences with Col-0 (Mann and Whitney test; α=5%).

**Fig. S10.**
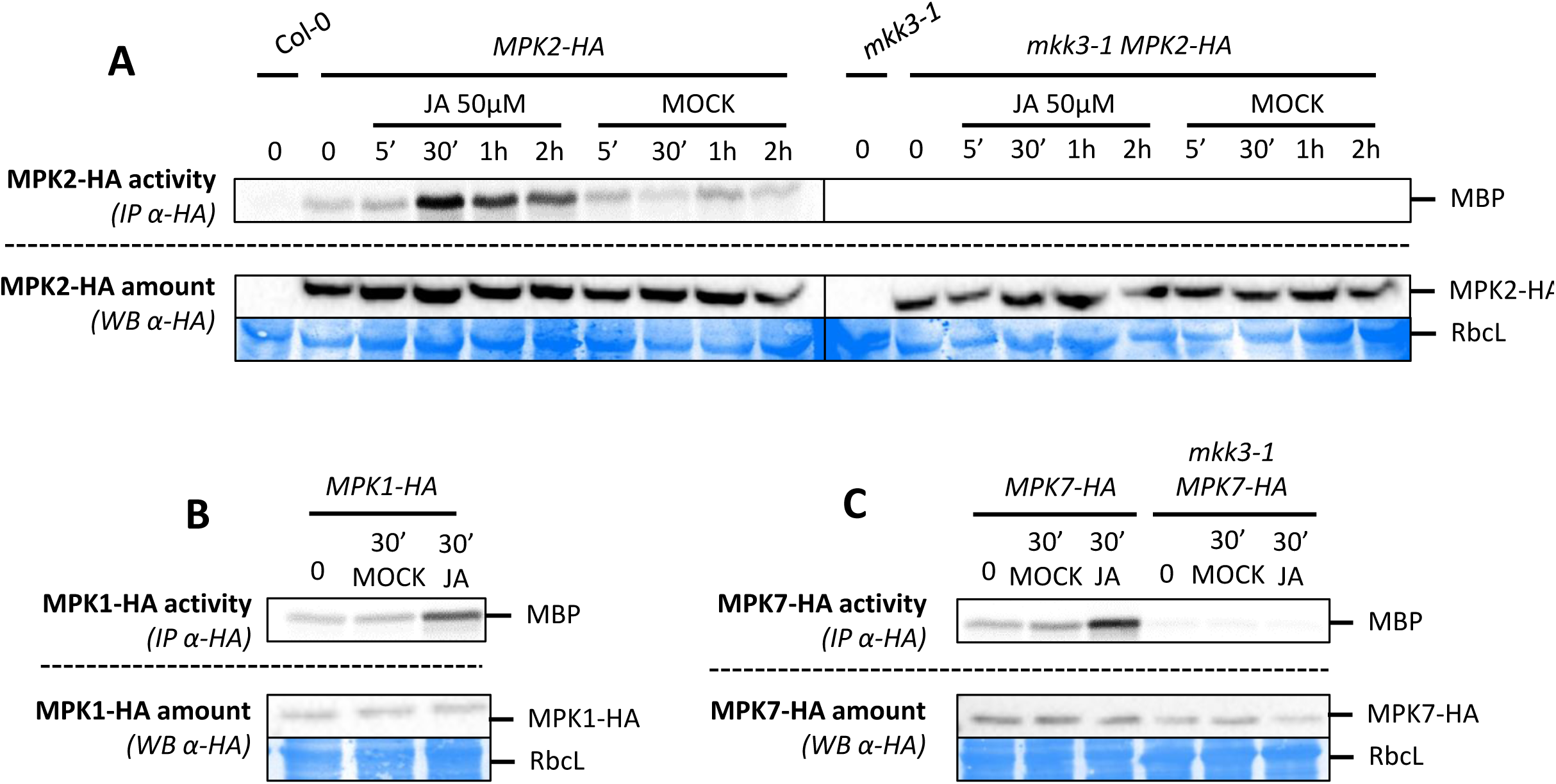
MPK1, MPK2 and MPK7 are activated by JA. **A to C.** Kinase activity of MPK2 (A), MPK1 (B) and MPK7 (C) after immunoprecipitation with an anti-HA antibody from leaves of WT (A, B, C) and *mkk3-1* (A, C) plants expressing a HA-tagged version of MPK2 (A), MPK1 (B) and MPK7 (C) following 50µM JA or MOCK (EtOH) treatments for indicated times. Protein amount is monitored by western-blot using anti-HA antibody. Equal loading is controlled by Coomassie staining.

**Fig. S11.**
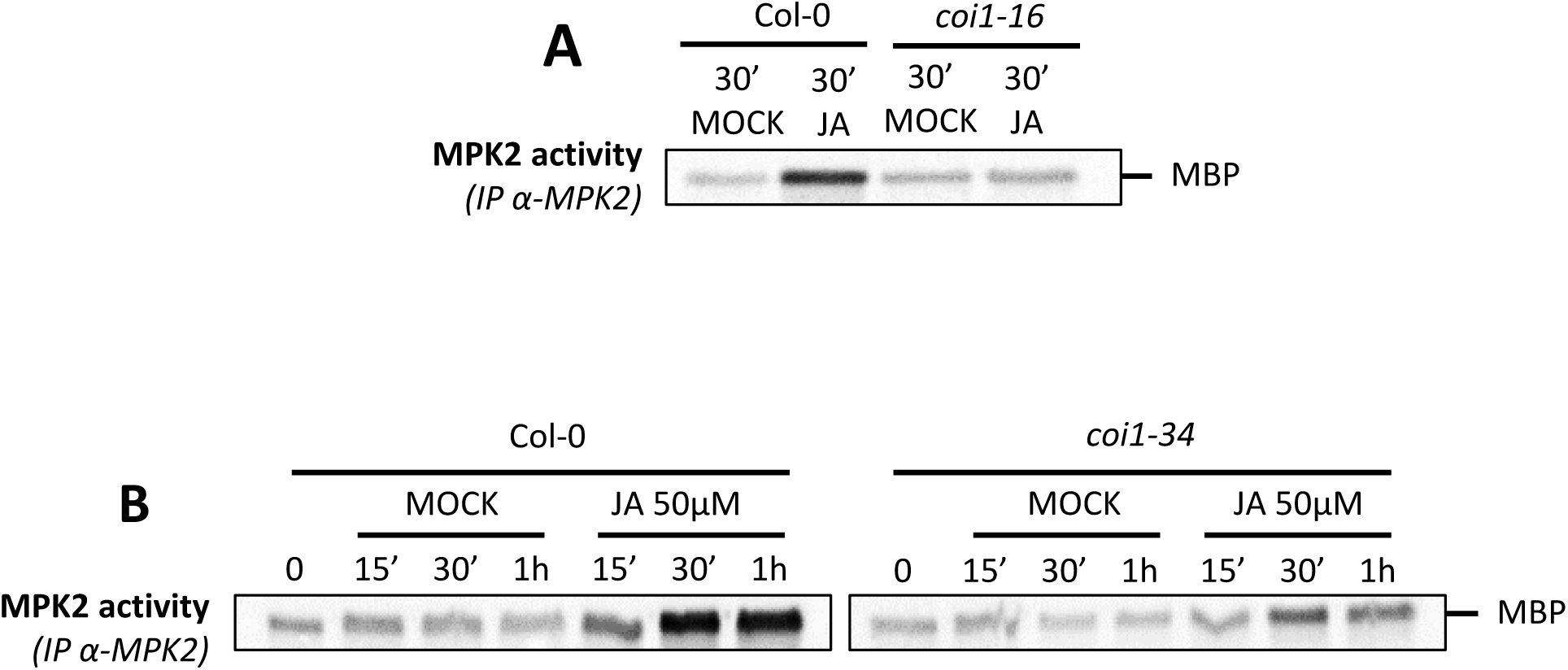
JA triggered MPK2 activation is impaired in mutants of JA receptor. **A and B.** Kinase activity of MPK2 after immunoprecipitation with an anti-MPK2 specific antibody from Col-0 (A, B) *coi1-16* (A) and *coi1-34* (B) plantlets following 50µM JA or MOCK (EtOH) treatments.

**Fig. S12.**
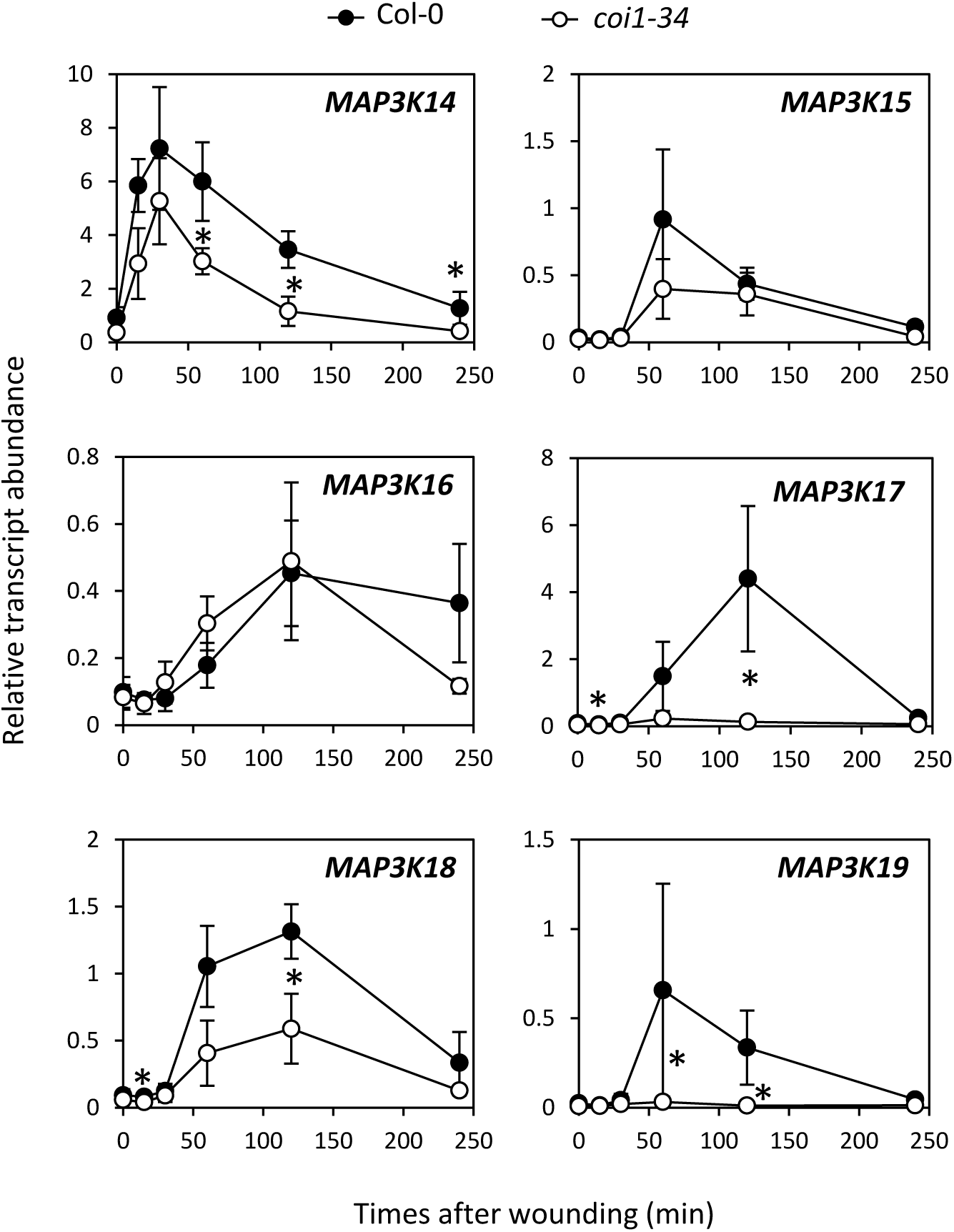
*coi1-34* mutation reduces the expression of some wounding-induced *MAP3Ks*. qRT-PCR analysis of clade-III *MAP3Ks* in response to wounding in Col-0 and *coi1-34* leaves. Transcript levels are expressed relative to *ACTIN2* reference gene. Values are mean±SE of 3 biological replicates. Stars indicate statistical differences between WT and *coi1-34* (Mann and Whitney test; α= 1%).

**Fig. S13.**
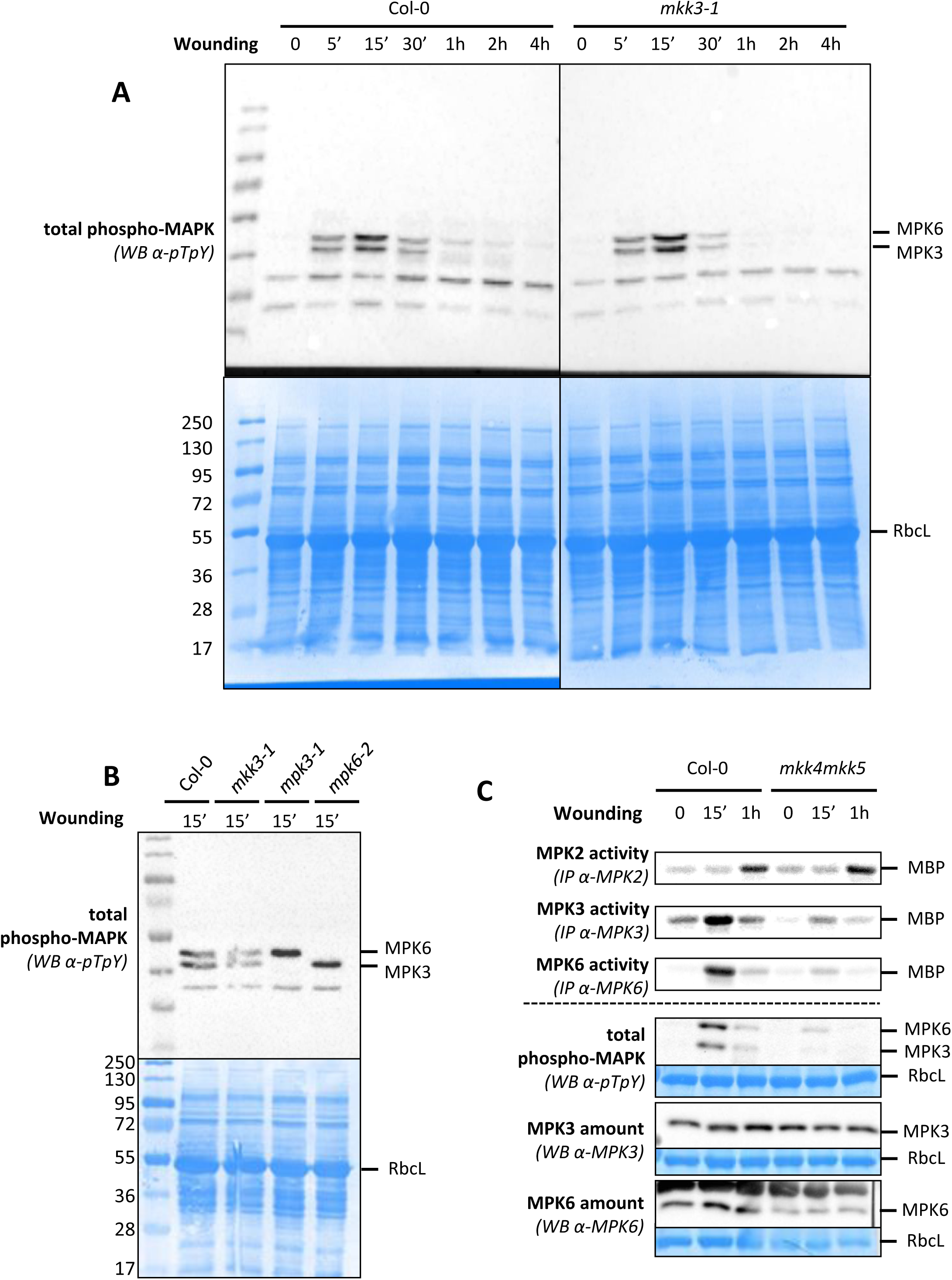
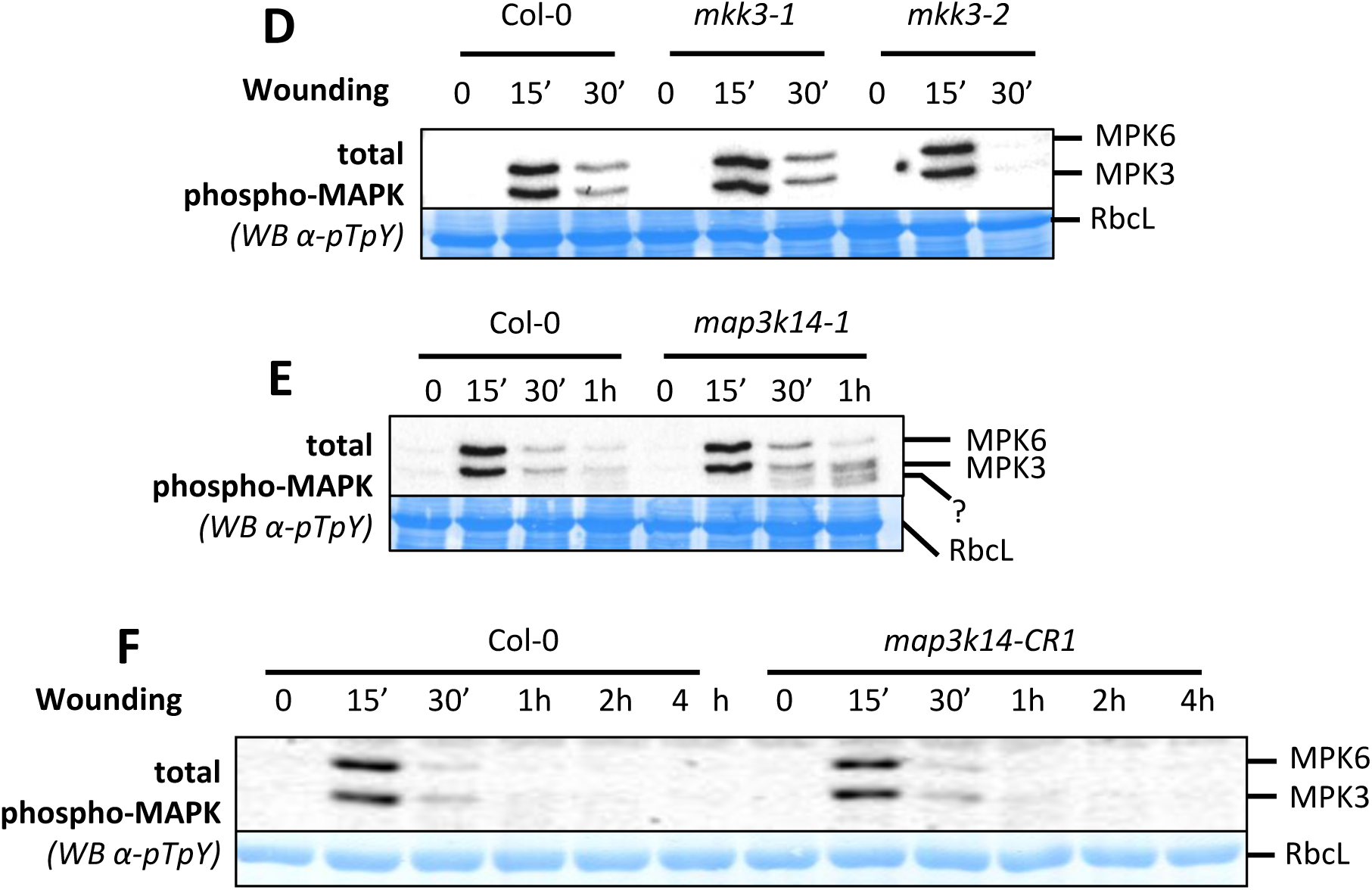
MPK3 and MPK6 activation depends on MKK4/MKK5 but not on MKK3. **A and B.** Western-blot using antibody raised against the phosphorylated form of ERK2 (anti-pTpY) in indicated genetic backgrounds after wounding. Equal loading is controlled by Coomassie staining. Figure S13A is an un-cropped version of western blot shown in figure 6A. **C.** Kinase activity of MPK2, MPK3, MPK6 after immunoprecipitation with an appropriate specific antibody from wounded leaves of Col-0 and *mkk4mkk5* plants. MPK3/6 activation was monitored by western-blot using antibody raised against the phosphorylated form of ERK2 (anti-pTpY). Equal loading is controlled by Coomassie staining. **D to F.** Western-blots using antibody raised against the phosphorylated form of ERK2 (anti-pTpY) in indicated genetic backgrounds. “?” (D) referred to a *map3k14-1* specific band which could correspond to the over activation of clade-C MAPKs. Equal loading is controlled by Coomassie staining.

**Fig. S14.**
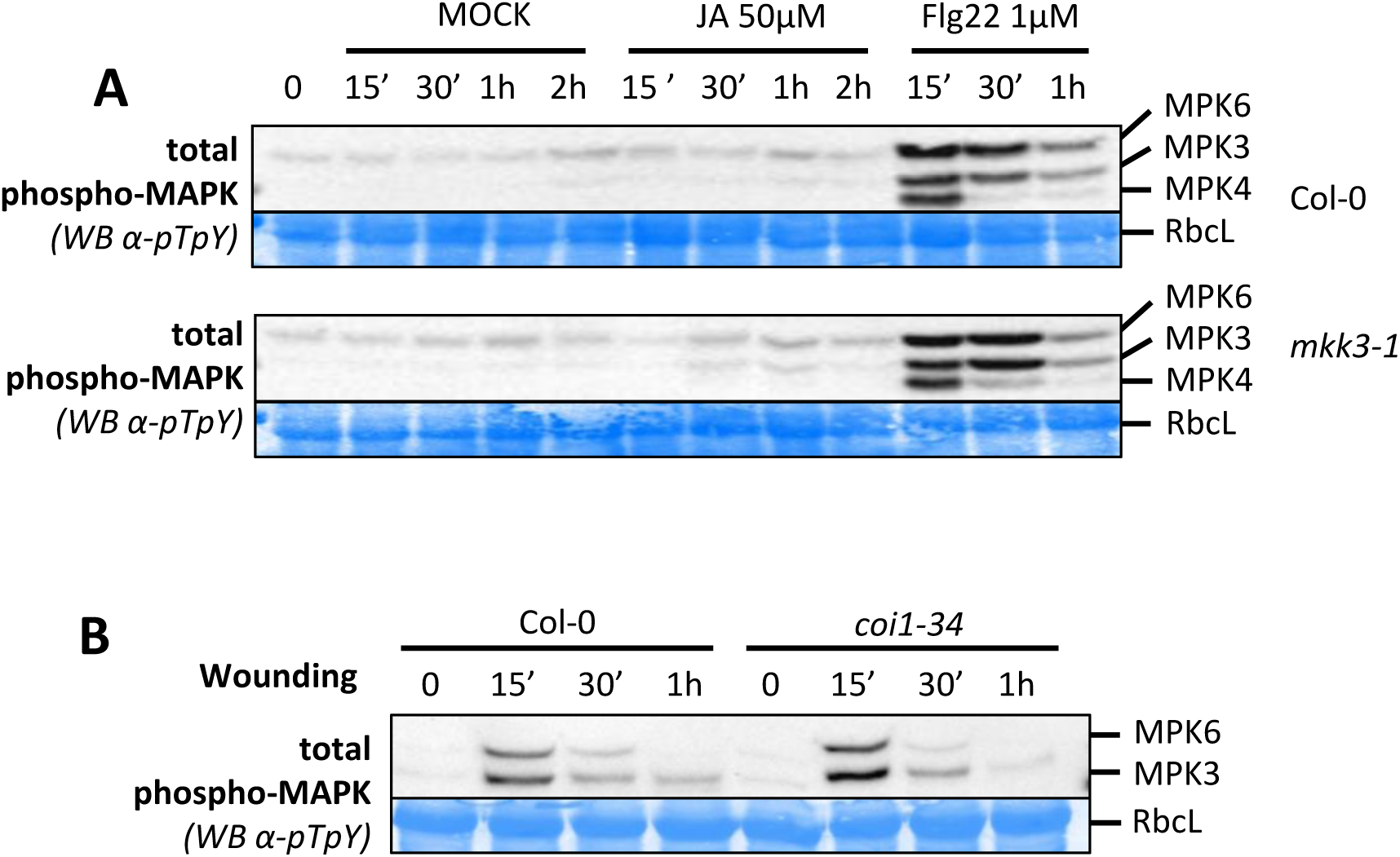
MPK3 and MPK6 activity is independent of JA. **A.** Western-blots using antibody raised against the phosphorylated form of ERK2 (anti-pTpY) in *in vitro* Col-0 and *mkk3-1* plantlets treated with 50µM JA, 1µM flg22 and MOCK (EtOH). Equal loading is controlled by Coomassie staining. **B.** Western-blots using antibody raised against the phosphorylated form of ERK2 (anti-pTpY) in Col-0 and *coi1-34* in response to wounding. Equal loading is controlled by Coomassie staining.

**Fig. S15.**
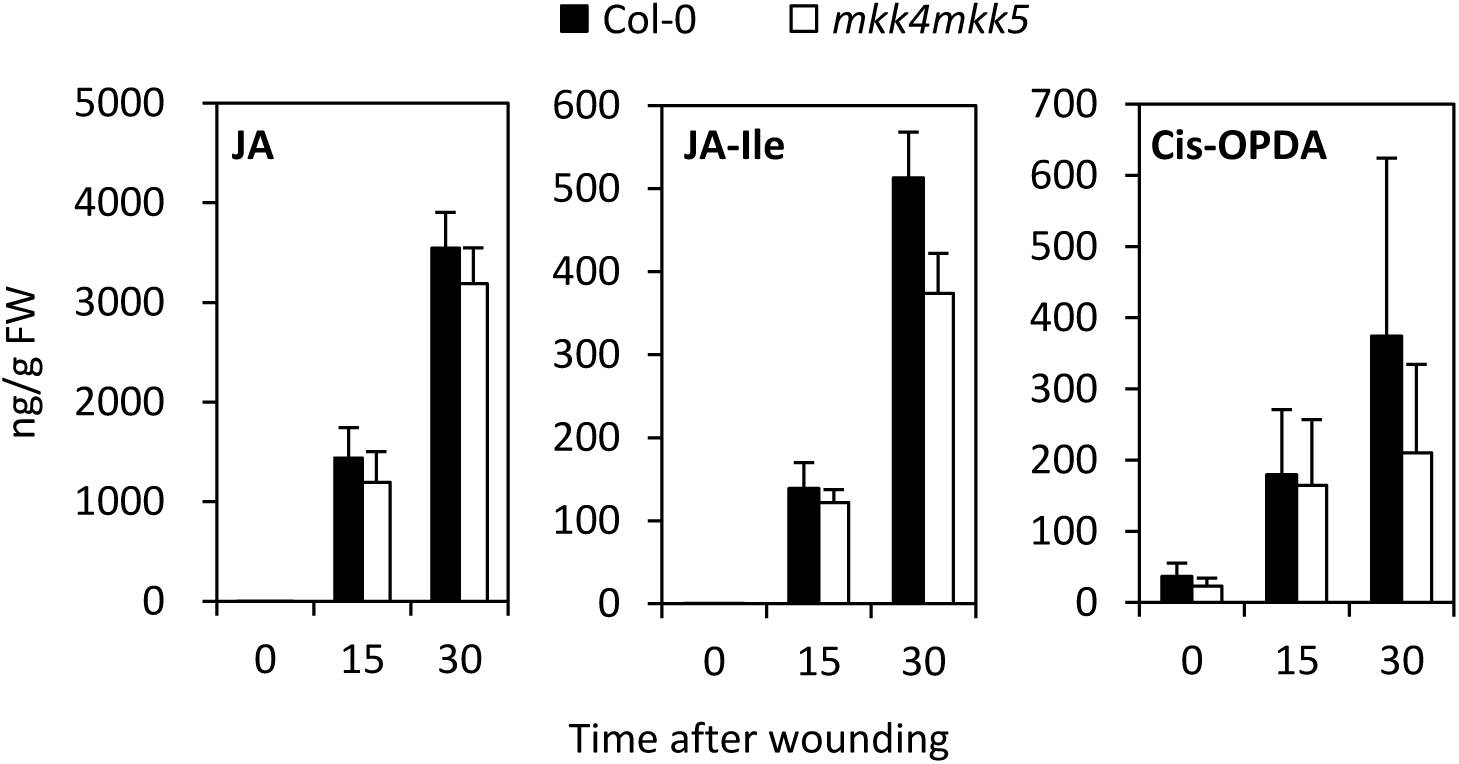
Oxylipin contents in *mkk4mkk5*. JA, JA-Isoleucine and Cis-OPDA contents in wounded leaves of Col-0 WT and *mkk4mkk5*. Values are mean±SE of 3 biological replicates.

**Fig. S16.**
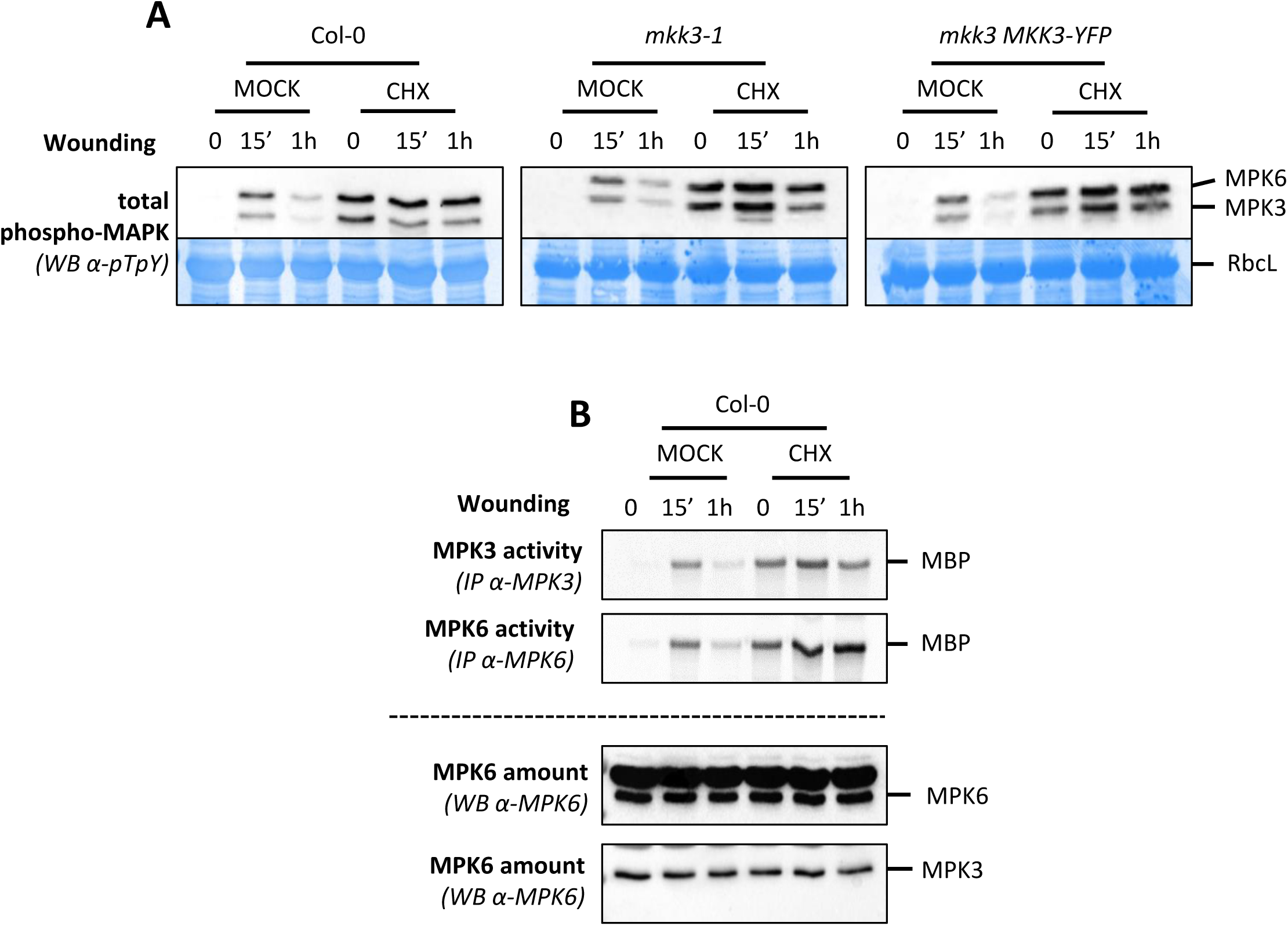
MPK3 and MPK6 are activated by cycloheximide. **A.** Western-blots using antibody raised against the phosphorylated form of ERK2 (anti-pTpY) from leaves of indicated genetic backgrounds following 100µM CHX and MOCK (DMSO) spraying prior to wounding. Equal loading is controlled by Coomassie staining. **B.** Kinase activity of MPK3 and MPK6 after immunoprecipitation with specific antibodies from Col-0 leaves following 100µM CHX and MOCK (DMSO) spraying prior to wounding. Western-blots show MAPK amount.

## Typical sample preparation and kinase assay to monitor kinase activity after wounding

### A. PREPARATION OF THE WOUND SAMPLES

#### 4-5 weeks before: preparation of plants

- Keep seeds in water at 4°C for 1-2 days to homogenize germination.
- Rehydrate the pellets in their holder (Jiffy-7 38mm Pellet-Pack, Ref # 32204011, Jiffy Products International AS, Norway) kept on a tray for 1h by bathing with tap water. Remove the extra water by draining it out of the tray.
- Identify the pellets which will receive the seeds of each genotype by writing on the plastic holder. Genotypes have to be randomized. Typically for two genotypes (A and B), we alternate the plants of each genotypes (see tray map).
- Sow 2-5 seeds per pellet.
- After 7-10 days, remove plantlets to keep a single healthy one per pellet.
- Grow plants during 4-5 weeks at 12h/12h light (µmol m-2s-1/dark, 70% humidity and 22°C. They are watered on Monday, Wednesday and Friday by filling the tray with 2 cm tap water. Extra water that has not been sucked by pellets is removed after 1 hour.

#### Wounding experiment (Example of the comparison of two genotypes (WT versus mutant) for MAPK activation at 0’, 15’, 30’, 1h, 2h and 4h)

- Prepare the appropriate number (12) of 2mL tubes, each containing a 4-5 mm iron bead. - In the morning, prepare the experiment, by deciding which plants will be in the samples (typical tray map provided as example, W=wounding). We use usually 3 plants per sample, each plant being wounded on 3 fully developed leaves.
- Following the time scale, wound 3 fully grown leaves per plant.
- Harvest rapidly the 9 leaves (3 plants × 3 leaves/plant) in 2mL tube and freeze them in liquid nitrogen.
- Keep at −80°C until use.

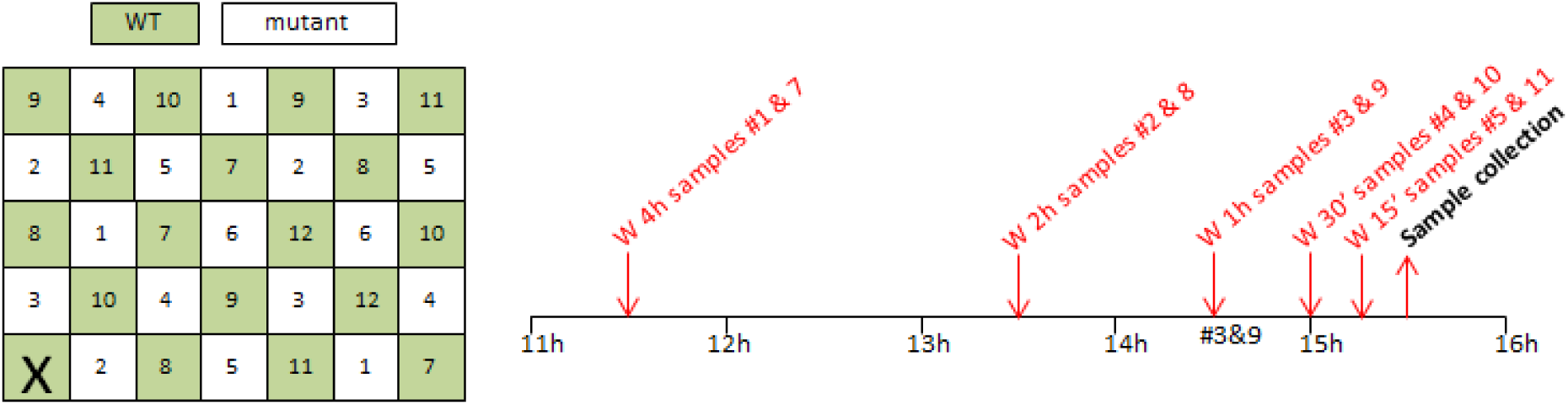

### B. KINASE ASSAY FOR PLANT CLADE-C MAPKS

**Adapted from Ortiz-Masia et al. (2007 FEBS letter) by Cécile Sözen and Jean Colcombet**

#### Day0 – Prepare day 1

- Label 7 sets of 1.5 mL tubes with sample number/names.
- Grind frozen samples either using iron beads/homogenizer. Keep at −80°C.

#### Day1 – Before to start

- Prepare the extraction buffer and cool on ice. For 100 mL **OB (**Ortiz-Masia Buffer) (25-30 samples):

10 mL Buffer E 10X (see stock preparation on the last page)
2 mL NaF 1M
2 mL β-glycerophosphate 1M
500 µL DTT 1M
2 tabs of EDTA-free protease inhibitors cocktail (Roche 04 693 132001)
- Cool down the bench microfuge at 4°C

#### Day1 – Extract soluble proteins

- Add 1 mL of OB to 300-350 mg of plant powder and keep on ice.
- Vortex several times to ensure mixing the powder and the buffer.
- Centrifuge 5 min at 14000 rpm at 4°C.
- Transfer on ice the supernatant to a new tube (set 1) and centrifuge again 5 min at 4°C.
- Transfer on ice the supernatant to a new tube (set 2) and centrifuge again 5 min at 4°C.
- The supernatant, which constitutes the protein extract, is transferred on ice to a new tube (set 3).

#### Day1 – Protein quantification using Bradford

- In new tubes (set 4) at RT, dilute your samples 20 times with OB.
- Fill the necessary wells of a 96 wells flat-bottom transparent plate with 250 µL Bradford reagent (Coomassie Protein Assay Kit, Thermo Scientific).
- All measurements are performed in triplicate. Drop 3x 10µL of the BSA standards in the left part of the plate from A1 to A3 (0.5 µg/10 µL) to F1 to F3 (2.5 µg/10 µL).
- Drop 3 × 10 µL of your samples starting from A4. Do not forget OB alone for reference.
- Wait few minutes before reading the results in the plate reader at 595nm.
- Determine protein concentration in each sample and calculate the dilution to obtain 0.5 µg.µL^−1^. In new tubes (set 5), prepare this dilution for each sample and keep on ice.

#### Day1 – Immunoprecipitation

- To prepare protein A-sepharose beads (Invitrogen, ref 101042), pipette 30 µL of the slurry per reaction in a 2 mL tube.
- Wash 3 times with 1 ml OB using the bench microfuge.
- Adjust the final volume of bead slurry with OB to get 60 µL per sample.
- Add the antibody to the bead slurry: 2 µL anti-MPK1/2/7 crude sera or 0.5 µL anti-HA (SIGMA H6908) per reaction
- Transfer 200 µL (100 µg of protein) of normalized samples from Set 5 to new tubes (Set 6). Keep on ice both set of tubes.
- Add 60 µL of bead-antibody slurry using a cut tip. Homogenize the slurry before each pipetting.
- Incubate the tubes on a wheel at 4°C for 2-3 hours.
- Transfer 50 µL of normalized samples (Set 5) to new tubes (Set 7). Add 50 µL Laemmli buffer 2X, heat at 95°C for 5 minutes and freeze for later western-blot.

#### Day1 - Washing

- Prepare 40 mL of kinase buffer

1.2 mL Tris-HCl pH7.5 1M (final 30 mM)
80 µL EGTA 0.5M (final 1 mM)
400 µL MgCl2 1M (final 10 mM)
40 µL DTT 1M (final 1 mM)
800 µL β-glycerophosphate 1M (final 20 mM)
- Centrifuge samples 20 sec at 10000 rpm at 4°C.
- Keep the tubes on ice. Discard the supernatant but leave 50 µL including beads in the bottom.
- Add 1 mL OB, mix by inverting, centrifuge 20 sec at 10000 rpm at 4°C and discard supernatant.
- Repeat 1 additional wash with OB then wash once with 1 mL kinase buffer.
- Keep the tubes on ice. Discard supernatant but leave 50 µL including beads in the bottom.
- Centrifuge again samples 20 sec at 10000 rpm at 4°C to pull down droplets.

#### Day1 – Kinase assay

- Keep tubes on ice while preparing the kinase reaction buffer. Premix per reaction:

15 µL Kinase Buffer
1.5 µL MBP 10mg/mL
0.15 µL ATP 10mM
- Use 200µL tips to remove the remaining supernatant from beads.
- In the radioactive area, transfer the sample tubes and kinase reaction premix in plexiglass holder at RT.
- Add the ^33^P [ATP] (2µCi per reaction) to the premix. Mix carefully by pipetting.
- Transfer 15 µL of radioactive kinase reaction buffer premix to each sample.
- Incubate for 30 minutes at RT.
- Stop the reaction by adding 15 µL Laemmli buffer 2X, mix carefully by pipetting and heat 5 minutes at 95°C.
- Freeze the samples

#### Day1 – Prepare day 2

- Prepare an appropriate number of SDS-PAGE gels (15% acrylamide, 1 mm thick) to run the kinase assay reaction. Keep them at 4°C overnight.
- If necessary, prepare an appropriate number of SDS-PAGE gels (10% acrylamide, 1 mm thick) to perform western-blot. Keep them at 4°C overnight.

#### Day2 – Kinase assay

- In the radioactive room, run the reaction samples (10 µl) on 15% SDS-PAGE until the migration front (which contains free radiolabelled nucleotides) runs out of the gel.
- Stain the gels using Coomassie solution for 1 h followed by washes in destaining buffer (2 h or over-night).
- Dry gels using a gel dryer and expose to a PhosphorScreen for 3 days. Detect radioactive bands with a Typhoon Imaging system (GE Healthcare).
- Take a picture of the gel for Coomassie loading control.

#### Preparation of the stock solutions

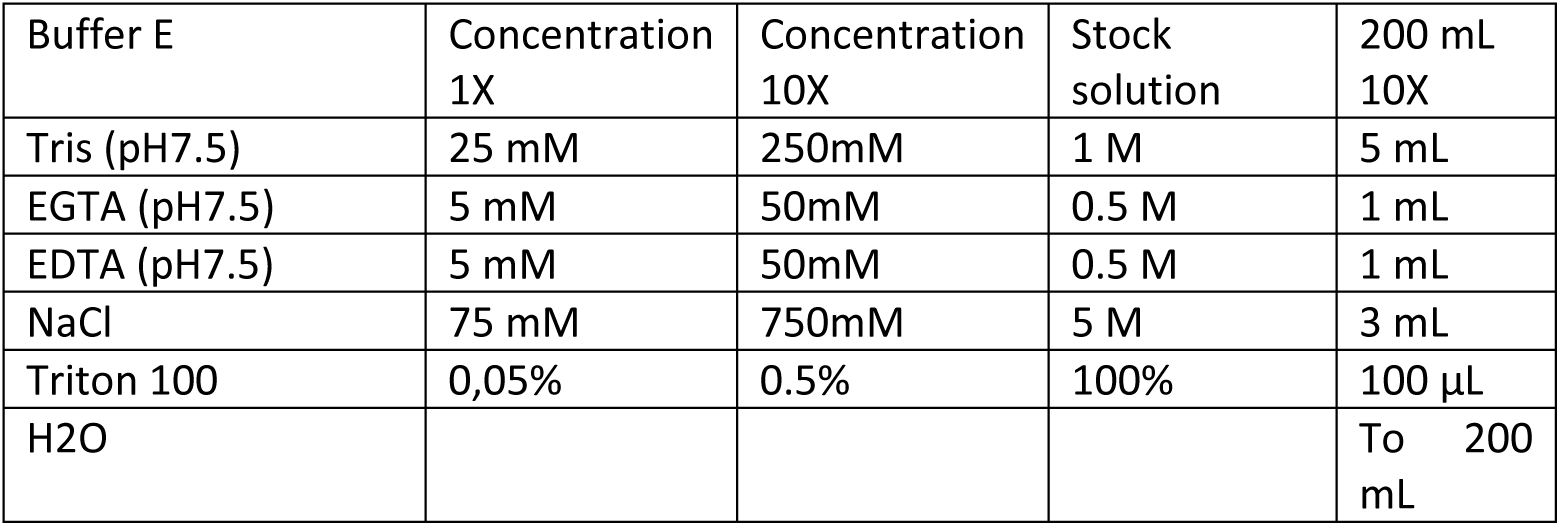

Prepare aliquots of 10mL and store at −20°C.

- NaF 1M:

M= 41.99g/mol
2.1g for 50mL.
Prepare aliquots of 2mL and store at −20°C.
- β-glycerophosphate 1M:

M= 216.04g/mol
10.8g for 50mL.
Prepare aliquots of 2mL and store at −20°C.
- DTT 1M:

M= 154.25
1.54g for 10mL.
Prepare aliquots of 500µL and store at −20°C.
- BSA range

From a BSA solution of 2µg/µl (Bovine Serum Albumin Standard, Thermo Scientific™, Ref: 23209)
Prepare a range from 0.5µg/10µl to 2.5 µg/10µL.
0.5µg/10µL = 25µL BSA 2µg/µL + 975µL H_2_O
0.75µg/10µL = 37.5µL BSA 2µg/µL + 962.5µL H_2_O
1µg/10µL = 50µL BSA 2µg/µL + 950µL H_2_O
1.5µg/10µL = 75µL BSA 2µg/µL + 925µL H_2_O
2µg/10µL = 100µL BSA 2µg/µL + 900µL H_2_O
2.5µg/10µL = 125µL BSA 2µg/µL + 875µL H_2_O
Prepare aliquots of 40µL and store at −20°C.
Do not freeze again once defrost!

---

**Supplemental table 1.**
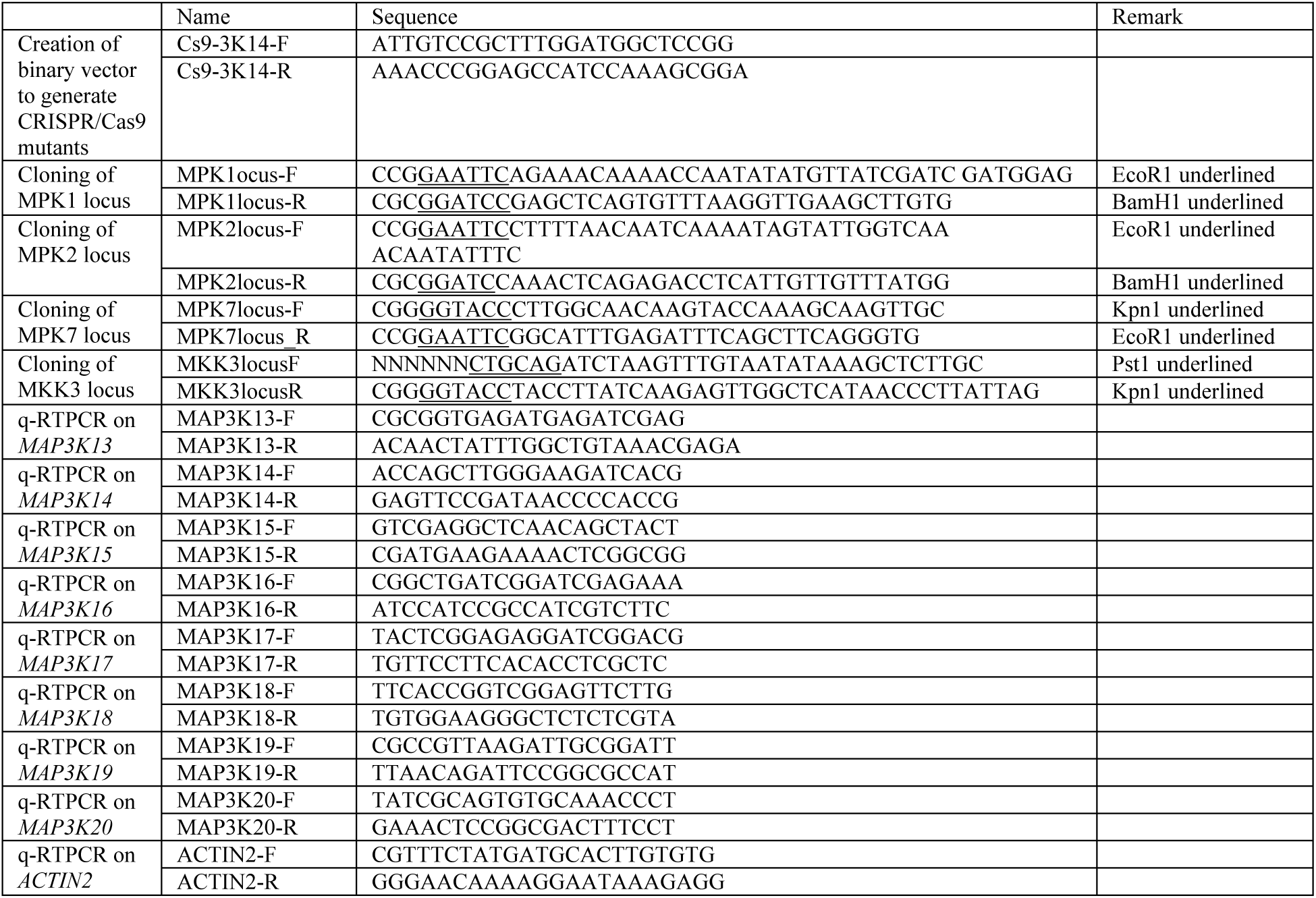
Primers used in this study.

**Supplemental table 2.**
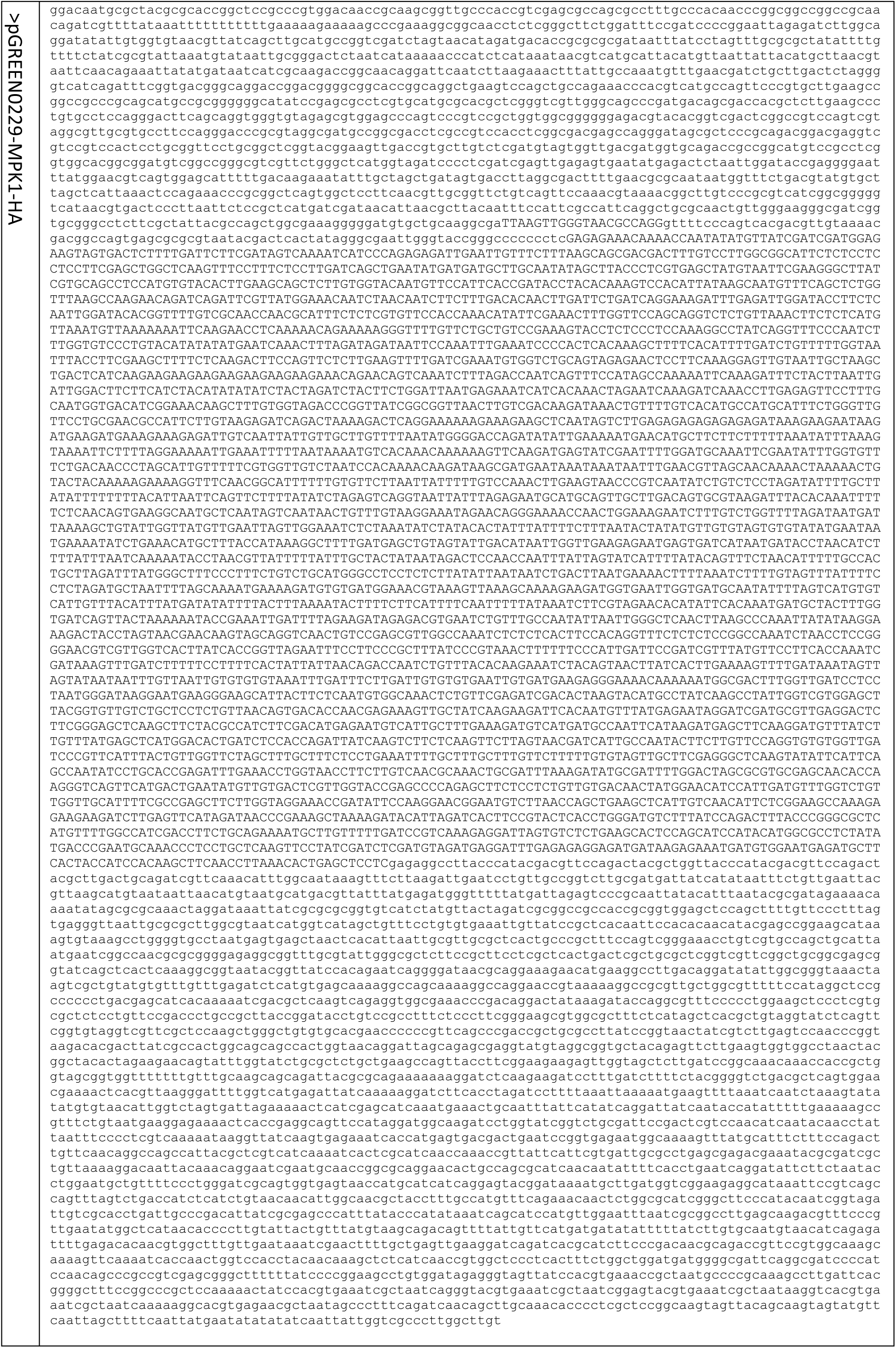
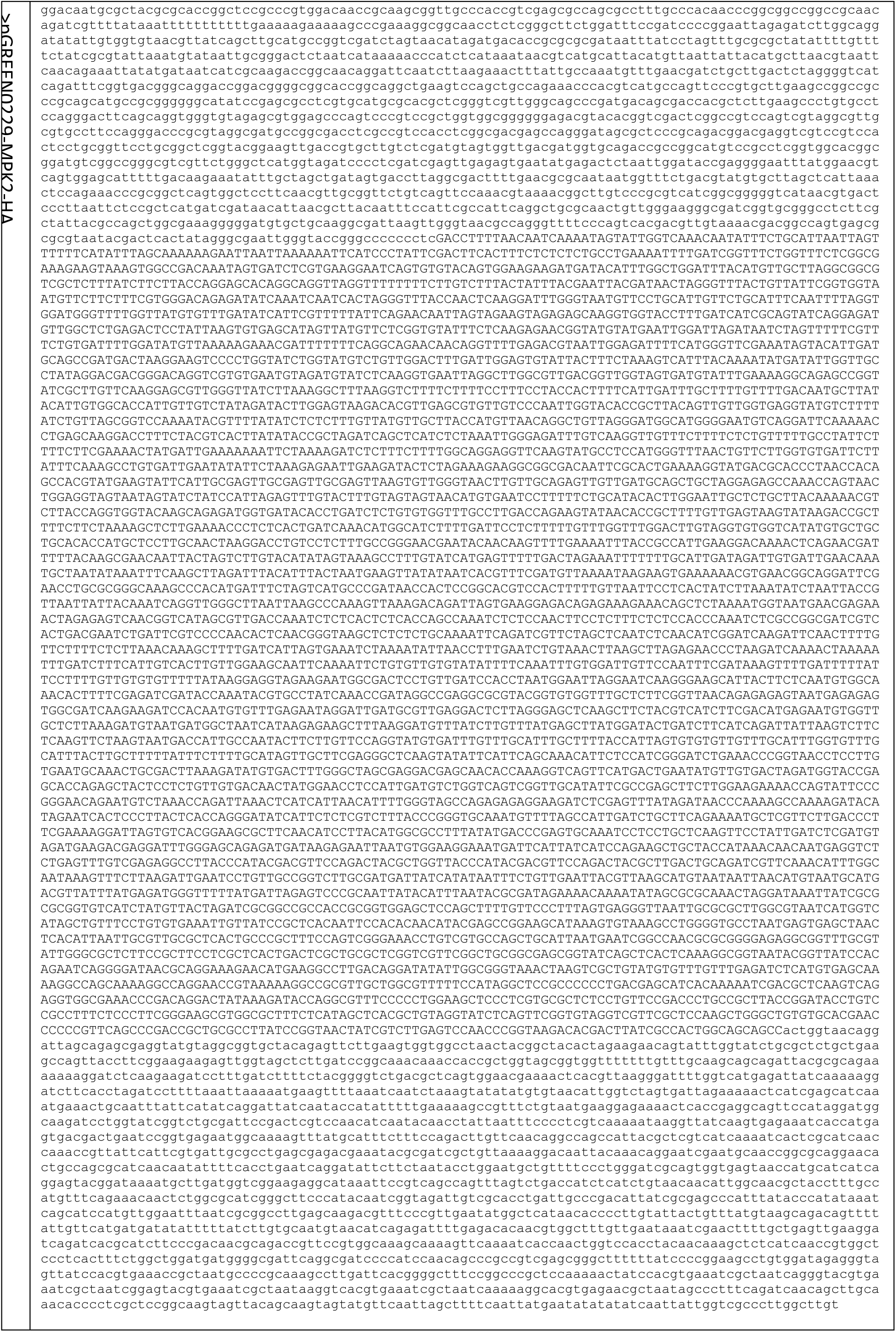
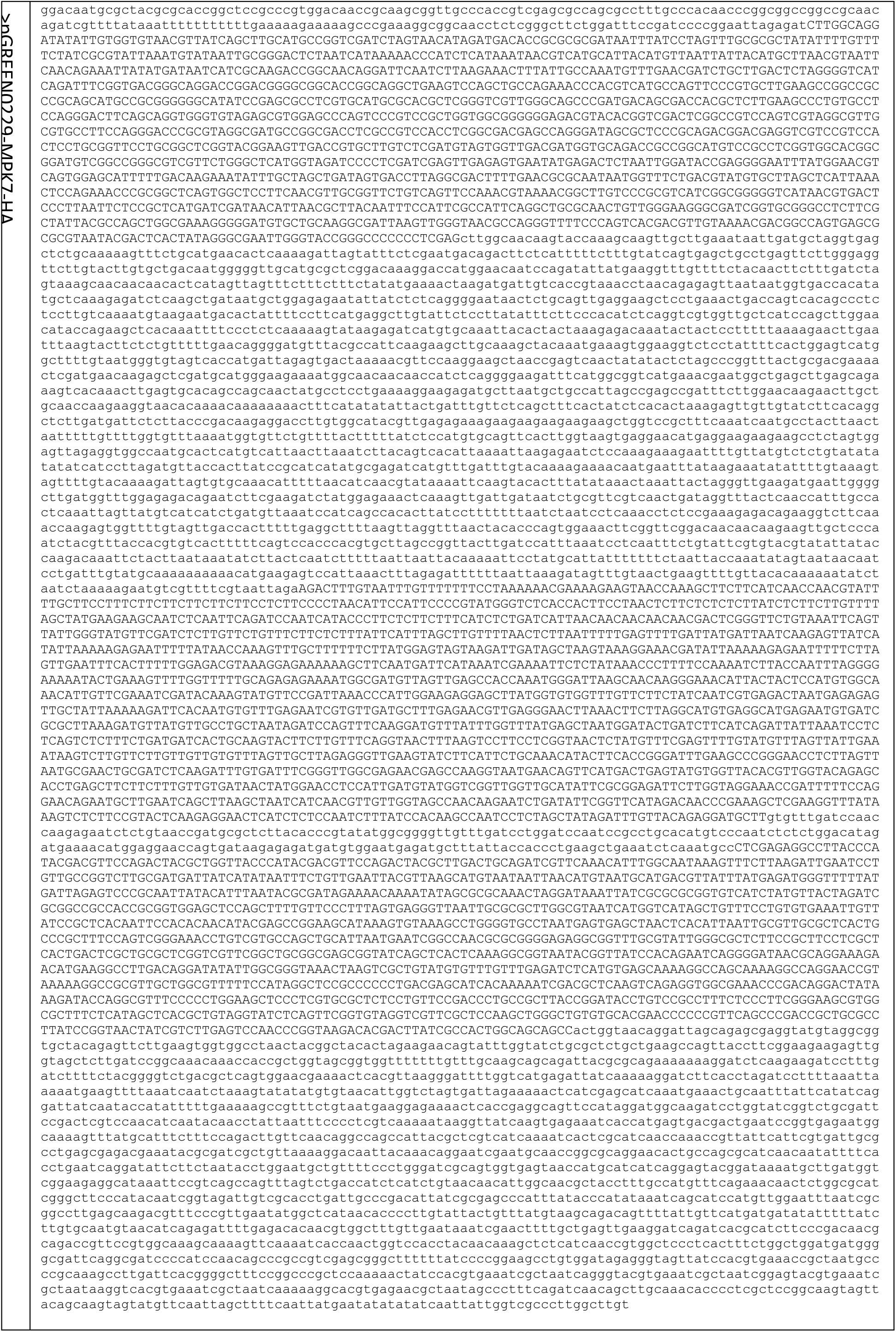
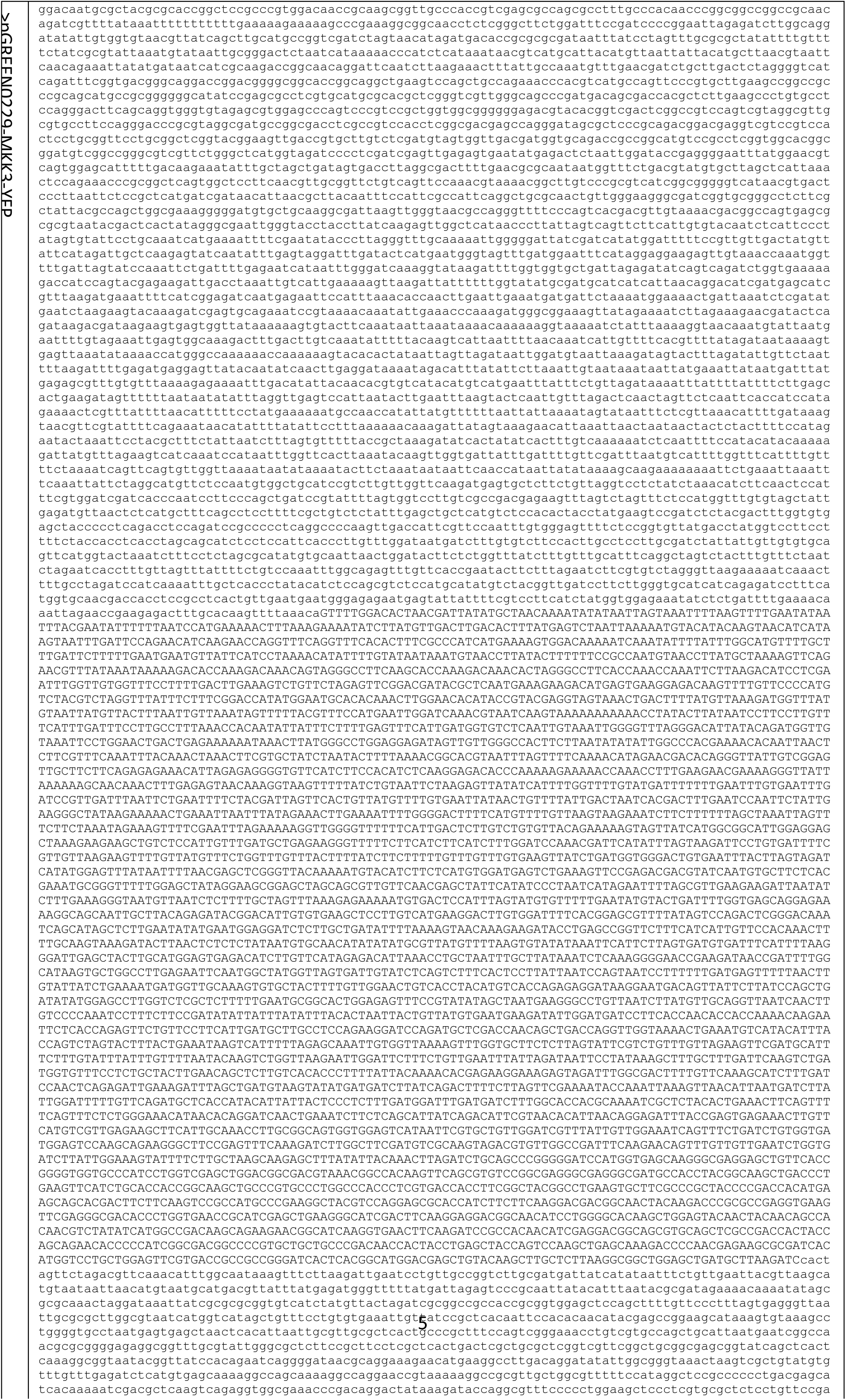
Sequences of plasmids used to generate complemented transgenic lines.

